# Mapping the Telomeric 3D Interactome with Telomere-C Reveals Repetitive Element Hubs Associated with Telomere Maintenance

**DOI:** 10.1101/2025.10.09.681516

**Authors:** Yi-An Chen, Ogechukwu Mbegbu, Noelle H. Fukushima, T. Rhyker Ranallo-Benavidez, Tianpeng Zhang, Floris P. Barthel

## Abstract

Telomeres are essential for genome integrity, but the accurate, high-resolution mapping of their three-dimensional (3D) chromatin interactions, a process thought to mediate gene regulation and telomere maintenance, has been limited by the repetitive nature of subtelomeric DNA and shortcomings of traditional sequencing-based chromatin conformation capture methodologies. To overcome these challenges, we developed Telomere Conformation Capture by sequencing (Telomere-C), a novel high-throughput approach for the genome-wide identification of telomeric chromatin interactions. We applied Telomere-C to three normal fibroblast cell lines (BJ, IMR90, WI38), one telomerase-positive cancer cell line (HeLa S3) and two cancer cell lines (U2-OS and WI38-VA13) that maintain telomeres by alternative lengthening of telomeres (ALT). We generated millions of telomere-associated reads per sample and identified up to 41,026 interaction peaks. We successfully validated previously-described telomeric interaction sites (e.g., *TERT* and *DUX4*), and critically, revealed a novel class of ultra-long-range interactions extending over 5 Mb away from the telomere, which surprisingly constitute over 80% of all identified contacts. Contrary to prior studies focusing on protein coding gene interactions, we discovered that the telomeric 3D interactome is overwhelmingly anchored at repetitive element hubs, particularly interstitial telomeric sequence (ITS), telomere-associated repeat 1 (TAR1) and D20S16 elements. We found that the clustering of these interactions correlates strongly with cell-type-specific telomere maintenance. Most notably, D20S16 telomeric interactions were uniquely and highly enriched in ALT cancer cells, suggesting a mechanistic link. Taken together, our study effectively constructs the first high resolution maps of the telomeric 3D interactome, redefining its scope to be dominated by ultra-long-range contacts with repetitive elements. This work provides fundamental insights into telomeric nuclear organization and establishes the telomeric D20S16 interaction as a molecular signature for the ALT pathway.

## Introduction

Chromatin organization plays a fundamental role in cell biology by organizing DNA and associated proteins in three-dimensional (3D) space. It is critically important for transcription, DNA replication and genome integrity by modulating the spatial proximity of coding and non-coding (regulatory) elements. Importantly, alterations in chromatin organization have been linked to human disease. For example, a recent study demonstrated that changes to chromatin accessibility in lamina-associated domains (LADs), caused by *LMNA* mutations, are associated with Hutchinson-Gilford progeria, a disease of premature aging linked to telomere dysfunction^1,2^.

Telomeres, repetitive DNA sequences at the ends of chromosomes, are fundamentally important for maintaining genome integrity^3^. Human telomere length ranges from 4-14 kb and terminates with a 130-210 bp single-stranded 3’ G-rich overhang^4–6^. Telomeric chromatin architecture is maintained by telomeric protein complexes, including shelterin, which plays a key role in differentiating telomeres from DNA double-strand breaks. Telomeres form unique chromosome-capping loop structures, called t-loops, by folding a single-stranded terminal overhang back onto upstream telomere sequences^7–10^. Previous studies showed that repression of the shelterin subunit TRF2 induces telomere deletions, resolution of t-loops, and chromosome fusions, highlighting the importance of telomeric chromatin organization in genome integrity^11,12^.

Beyond its role in end protection, telomeric chromatin was shown to epigenetically modulate gene expression via telomere position effects (TPE)^13^. In *Saccharomyces cerevisiae,* TPE is characterized by a transcriptionally silenced heterochromatin domain stretching from 2 - 20 kilobases away from the end of the chromosome^13–16^. The degree of heterochromatin spreading and consequential gene silencing was found to be associated with telomere length^13^, suggesting that telomeres form kilobase-scale loop structures to extend the range of TPE^14,17–19^. Several studies have also reported TPE “over long distances” (TPE-OLD) in humans and demonstrated telomere-length-dependent regulation of gene expression, with promoters up to 7.5 megabases away from chromosome ends^20–25^.

Telomeric chromatin organization is also important for telomere maintenance. For example, the alternative lengthening of telomeres (ALT) pathway exploits homologous recombination (HR), specifically break-induced replication, for telomere extension and is utilized by approximately 15%-20% of cancers^26–29^. Spatial proximity was shown to be an important mediator of recombination outcomes and is therefore an understudied variable governing the genetic consequences of ALT^30,31^. Recent studies have identified that orphan nuclear receptors (NRs), including COUP-TF1, COUP-TF2, TR2, and TR4, can bind to telomere variant (TCAGGG) repeats that are abundant in ALT-positive cells^32,33^. This binding allows the NRs to serve as a molecular scaffold for HR-based recombination^30,34^. Another study demonstrated that tethering of COUP-TF1 or COUP-TF2 to telomeres is sufficient to induce ALT phenotypes in normal human cells, highlighting the critical role of these NRs in ALT development by influencing telomeric chromatin organization^35^.

However, accurate detection and quantification of telomeric chromatin interactions remains challenging due to the repetitive and polymorphic nature of (sub)telomeric DNA. While image-based approaches are frequently used for validating telomeric chromatin interactions^36^, their low resolution and throughput have been bottlenecks in drawing robust mechanistic conclusions. Moreover, chromatin conformation capture sequencing approaches such as Hi-C^37^, ChIA-PET^38^, or HiChIP^39^ rely on enzymatic digestion that is incapable of fragmenting telomeric repeats and therefore falls short of capturing telomeric interactions^40^. Consequently, the long-range 3D interactome of telomeres, especially those governing pathways like ALT, remains largely unmapped.

Here, we developed a high-throughput sequencing approach, Telomere Conformation Capture by sequencing (Telomere-C), and mapped telomeric chromatin interactions in diploid human fibroblast cells, telomerase-positive cancer cells, and ALT-positive cancer cells. We identified a novel class of ultra-long-range interactions, which were found more than 5 Mb away from the telomeres. Unexpectedly, we found that telomeric interactions were highly enriched at repetitive elements, in particular interstitial telomeric sequence (ITS), telomere-associated repeat 1 (TAR1), and the poorly understood D20S16 repeat element. Traditional chromatin conformation capture (3C), Southern blotting and fluorescence *in situ* hybridization (FISH) validated these interactions. While most interactions were universally observed, we noted a particular enrichment of D20S16 element interactions in ALT cancer cell lines. These results provide critical new insights into telomeric chromatin interactions, demonstrating a novel class of ultra-long-range interactions and, notably, identifying D20S16 interactions as a specific, novel biomarker and potential mechanistic driver for ALT cancers.

## Results

### Telomere-C Reveals a Genome-wide Network of Ultra-Long-Range Telomeric Chromatin Interactions

In order to investigate how telomeres are organized in the nucleus, we developed Telomere-C to study telomeric chromatin interactions in cells in an unbiased manner (Figure 1A, Supplementary Figure 1A-C, and *Methods*). Briefly, chromatin from crosslinked cells was sheared and a telomeric probe was used to co-capture telomeric DNA and interacting DNA. Chromatin was de-crosslinked and probe-bound telomeric DNA was discarded, leaving the captured telomere-interacting DNA for sequencing. To study telomeric chromatin interactions in normal and cancer cells with varying telomere maintenance mechanisms, we applied Telomere-C to three normal fibroblast cell lines (BJ, IMR90, WI38), one telomerase-positive cancer cell line (HeLa S3) and two cancer cell lines (U2-OS and WI38-VA13) that maintain telomeres via ALT (Figure 1B). Sequencing reads were processed with our custom pipeline (Supplementary Methods and Supplementary Figure 1C), resulting in the identification of hundreds to thousands of Telomere-C peaks per library (Supplementary Figure 1D, Table 1).

**Figure 1.**
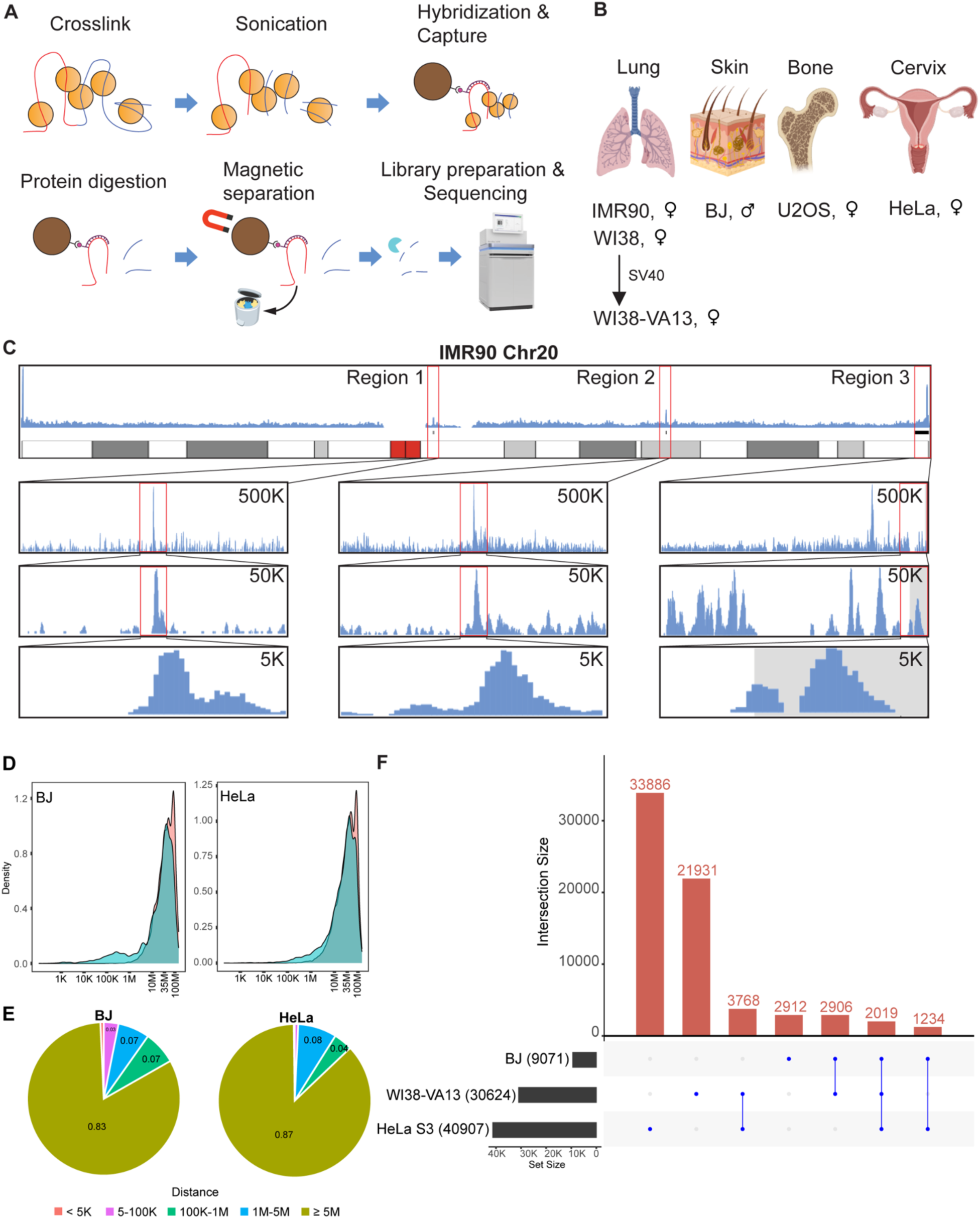
Detection of distal and proximal telomeric chromatin interaction using Telomere-C in normal, transformed, and cancer cell lines. **A** Schematic of the Telomere-C library preparation, including the enrichment of telomere-associated DNA. Yellow spheres, red threads, and blue threads represent chromatin-associated proteins, telomeric DNA, and telomere-associated DNA in chromatin, respectively. Chromatin was hybridized with a biotinylated telomere-targeting PNA probe (purple thread with a magenta sphere) followed by magnetic separation using streptavidin beads (brown spheres). Telomeric DNA bound to the beads was discarded, while the remaining telomere-associated DNA was digested with AluI (cyan 3/4 circle) and subsequently processed for Illumina library preparation and sequencing. **B** Source and sex of human normal, SV40-transformed, and cancer cell lines used for Telomere-C. **C** Normalized Telomere-C reads aligned to chromosome 20 from IMR90. The top panel shows an overview of the entire chromosome. Regions of interest are highlighted in rectangles, with three magnified windows: 500, 50, and 5 Kbp. The grey shading indicates the telomeric region. **D** Distribution of the relative distances of Telomere-C peaks to telomeres (cyan) or centromeres (coral) in the indicated cells. **E** Proportion of Telomere-C peak distances to telomeres. Distances are categorized by range: <5 Kb (coral), 5-100 Kb (purple), 100 Kb-1 Mb (blue), 1-5 Mb (green), and ≥ 5 Mb (olive) in the indicated cells. **F** An UpSet plot showing unique and shared Telomere-C peaks identified among the BJ, WI38-VA13 and HeLa S3 cells.

**Table 1.** Summary of Telomere-C peak sizes.

|  | Total peaks | Min | Q1 | Median | Mean | Q3 | Max |
| --- | --- | --- | --- | --- | --- | --- | --- |
| BJ | 9,397 | 31 bp | 302 bp | 602 bp | 626.6 bp | 902 bp | 12,602 bp |
| IMR90 | 1,257 | 31 bp | 302 bp | 602 bp | 868.9 bp | 902 bp | 11,102 bp |
| WI38 | 161 | 31 bp | 302 bp | 902 bp | 939.4 bp | 1202 bp | 5,402 bp |
| WI38-VA13 | 31,072 | 31 bp | 302 bp | 602 bp | 733.1 bp | 902 bp | 9,902 bp |
| U2-OS | 6,158 | 31 bp | 302 bp | 602 bp | 674.4 bp | 902 bp | 6,602 bp |
| HeLa S3 | 41,026 | 31 bp | 602 bp | 1202 bp | 1,184 bp | 1,502 bp | 25,502 bp |

To explore the distribution of telomeric chromatin interactions across a chromosome, we plotted normalized Telomere-C reads (see Methods) on chromosome 20 in IMR90 cells as a representative example and visually noted several distinct peaks (Figure 1C). Specifically, we observed peaks at the terminal region of each chromosome arm, a peak near the centromere and a peak at an intra-chromosomal locus on chromosome 20. Applying a custom peak calling algorithm (see Methods) and plotting these peaks in all tested cell lines showed a diffuse distribution of peaks at both telomeric and intra-chromosomal sites (Supplementary Figure 2A).

We hypothesized that telomeres are more likely to interact with elements that are closer in linear space compared to elements that are further apart. To test this hypothesis, we examined the distribution of Telomere-C peaks as a function of their distance to the telomere or centromere. This analysis revealed three distinct categories of interactions that varied in their distance to the telomere. The first two categories showed average distances of 1 Kb and 100 Kb, consistent with the previously described TPE and TPE-OLD interactions^16,20–22,24^. Unexpectedly, we identified a significant number of ultra-long-range telomeric chromatin interactions occurring at distances over 5 Mb away from the telomere (Figure 1D and Supplementary Figure 1B). Specifically, in BJ and HeLa S3 cells, 83% and 87% of telomeric interactions involved interaction partners over 5 Mb away from the telomere, respectively; whereas 14% and 12% of these interactions occurred between 100 Kb and 5 Mb. Finally, less than 5% of telomeric interactions were found within 100 Kb of the chromosome ends (Figure 1E). These results suggest that telomeres predominantly interact with distal genomic regions rather than proximal peri-telomeric sites. This finding was consistent in all tested cell lines (Supplementary Figure 2 B-C).

To evaluate whether Telomere-C peaks were shared between cell lines and types, we analyzed peak locations and quantified the number of overlapping and non-overlapping peaks. Amongst the three cell lines with the highest numbers of Telomere-C peaks, we found that *n* = 2,912/9,071 (32.1%) peaks were unique to BJ, *n* = 21,931/30,624 (71.6%) peaks were unique to WI38-VA13 and *n* = 33,886/40,907 (82.8%) peaks were unique to HeLa S3, indicating a dominance of cell-line specific telomeric chromatin interactions (Figure 1F). Nevertheless, we found that 2,019 peaks were shared between these three lines, possibly reflecting conserved sites important for maintaining nuclear organization. Intriguingly, *n* = 1,195 peaks were shared between the ALT cell lines U2-OS and WI38-VA13, implicating a common telomeric chromatin organization in cells showing this phenotype (Supplementary Figure 2D).

These results demonstrate that Telomere-C successfully detects genome-wide telomeric chromatin interactions, revealing a novel class of ultra-long-range interactions. Moreover, both shared and unique peaks were detected in our dataset, reflecting the cell-specific and common chromatin organization among tested cell lines.

### Telomere-C Outperforms Micro-C and Hi-C in Detecting Telomeric Chromatin Interactions

To validate our method’s superiority in highly repetitive genomic regions, we directly compared the performance of Telomere-C in identifying telomeric chromatin interactions to that of Micro-C and Hi-C in IMR90 cells. While restriction enzyme-based approaches like Hi-C are incapable of digesting telomeric repeats, Micro-C utilizes micrococcal nuclease (MNase) to allow for sequence-unbiased digestion, enabling it to incorporate telomeric fragments more effectively^41^.

We applied an in-house pipeline (see Methods) to select read pairs associated with telomeres, or specifically reads containing (TTAGGG)_10_, from Hi-C and Micro-C datasets. Telomere-C unequivocally outperformed both methods, identifying 10,987,734 telomere-associated reads compared to only 16,049 and 1,305 telomere-associated reads using Micro-C and Hi-C, respectively (Table 2). Due to these low read counts, we could not detect any peaks in the Hi-C or Micro-C data sets. In contrast, Telomere-C detected 1,257 robust peaks (Table 1).

**Table 2.** Comparison of Hi-C and Micro-C Technologies with Telomere-C. Reads in Micro-C and Hi-C were categorized based on the presence of at least 10 (TTAGGG) repeats in both reads (2R-map) or one read (1R-map)

|  | Telomere-C | Micro-C | Hi-C |
| --- | --- | --- | --- |
| Sequencing reads | Input: 17.8M<br>Capture: 40.4M | 101.2 M | 281.3 M |
| Filtered reads | Input: 12.4 M<br>Capture: 11.0M | 2R-map: 3.4 K<br>1R-map: 12.7K | 2R-map: 173<br>1R-map: 1.1K |
| Telomere associated reads | 10,987,734 | 16,049 | 1,305 |
| Called peaks | 1,257 | N.A. | N.A. |
| Percent of peaks with<br>≥1 intersecting read | 1,257 (100%) | 332 (26%) | 37 (3%) |

Despite their quantitative differences, 26% and 3% of Telomere-C peaks overlapped with reads identified by Micro-C and Hi-C, respectively. This indicates that Telomere-C accurately captures true long-range telomeric interactions while operating at a superior efficiency than other methods. Furthermore, Telomere-C captured over 10.9 million telomeric chromatin interactions (61.73% of reads) compared to 16.0 thousand interactions in Micro-C (0.02% of reads) and 1.3 thousand (< 0.01% of reads) in Hi-C.

To further validate the accuracy of Telomere-C calls, we visualized Telomere-C together with Micro-C signal in a selected region. Although Micro-C data was much sparser, Telomere-C peaks and Micro-C signal largely agreed (Figure 2A). Taken together, our data demonstrate that Telomere-C is a powerful approach for detecting telomeric chromatin interactions, easily outperforming Micro-C and Hi-C in both the number and range of robust interactions detected across the genome.

**Figure 2.**
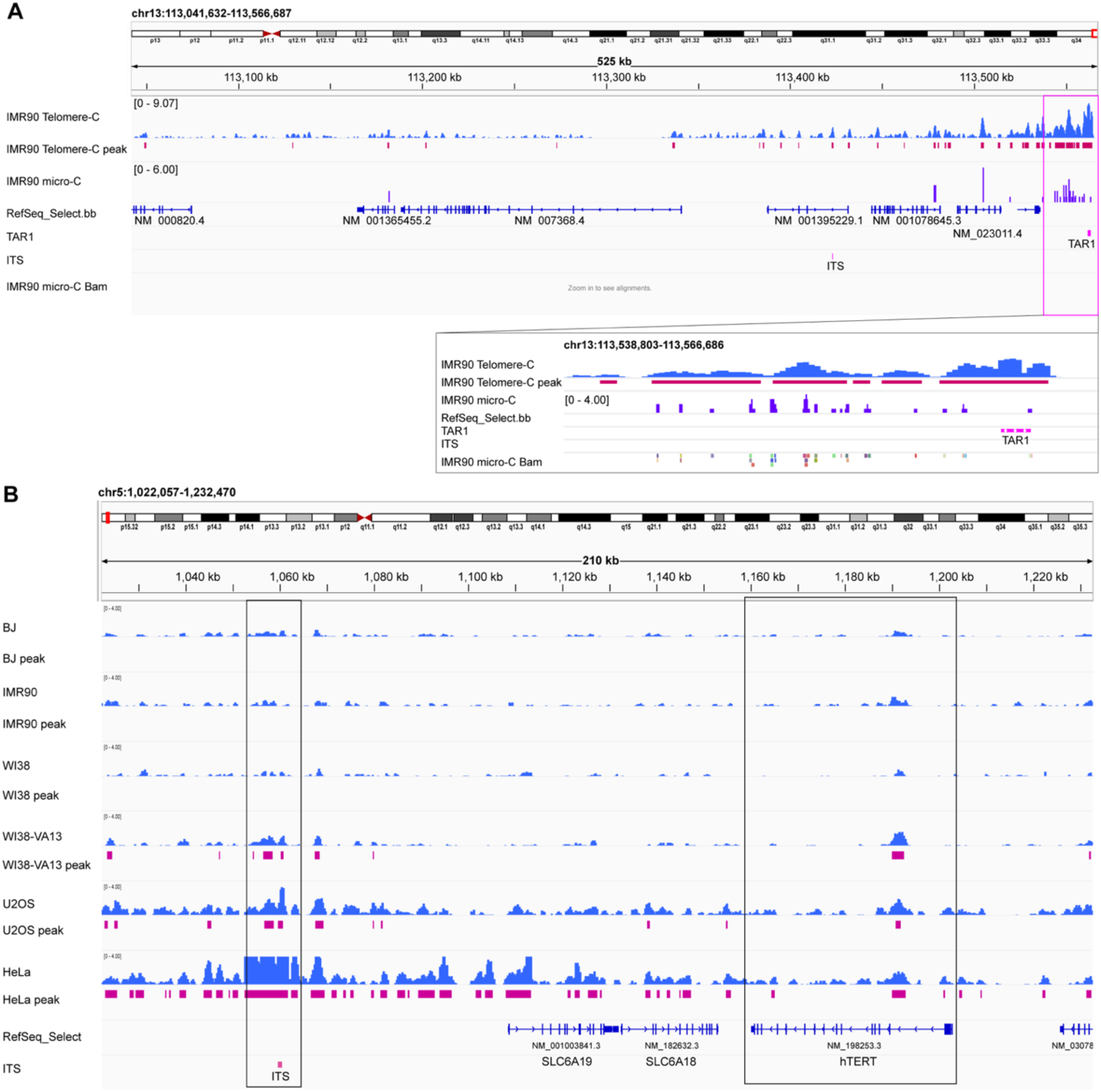
Validation of telomeric chromatin interaction in subtelomeric regions and at the *hTERT* locus. **A** The panel shows Telomere-C (blue bars) and Micro-C (purple bars) signals. Called Telomere-C peaks and repeat elements (ITS and TAR1) are shown as red and magenta solid rectangles, respectively. The magenta rectangle highlights a zoom-in window (chr13:113,538,803-113,566,686) showing raw sequence reads from IMR90 Micro-C data. **B** The panel shows Telomere-C signals (blue bars) and corresponding called peaks (red rectangles) for each indicated cell line. Two black rectangles highlight the *hTERT* transcript and the ITS locus 100 Kb downstream.

### Telomere-C Validates Known TPE-OLD Sites and Pinpoints Novel Interaction Loci

To verify that Telomere-C accurately captures established biological interactions, we sought to validate known long-range telomeric chromatin interactions previously implicated in TPE-OLD^22^. We focused on three genes: *TERT*, *DUX4* and *ISG15*.

First, we analyzed the TERT locus. We identified significant telomeric interactions adjacent to exon 2 of *hTERT* in both aneuploid (WI38-VA13, U2-OS, and HeLa S3) and normal cell lines (BJ, IMR90, and WI38), although the signal was much stronger in cancer cells (Figure 2B). Importantly, Telomere-C resolved a novel interaction site: an interstitial telomere sequence (ITS) located 100 Kb downstream of *TERT*. This locus had been previously proposed as a site of telomeric chromatin interaction but never validated at high resolution^22^.

Next, we analyzed the gene *DUX4*. Regulated by TPE, *DUX4* alternative splicing leads to the expression of the full-length toxic isoform DUX4-FL, which drives pathology in facioscapulohumeral muscular dystrophy (FSHD) patients^24^. Our Telomere-C data confirm a significant telomeric chromatin interaction 6 Kb downstream of *DUX4* in both cancer and normal cells (Supplementary Figure 3A).

Finally, we examined the *ISG15* locus. *ISG15* was previously found to be the target of a long-range telomeric chromatin interaction, but the precise interacting locus (which was known only to be within 200 Kb) was not known^25^. Telomere-C successfully mapped this interaction to exon 2 (BJ cells) and 5 Kb downstream of *ISG15* (BJ, IMR90 and WI38-VA13; Supplementary Figure 3B).

In summary, Telomere-C successfully validated multiple sites previously implicated as telomeric chromatin interactions. Moreover, owing to its higher resolution, our approach moves beyond the capacity of lower-resolution methods like FISH, enabling us to precisely pinpoint interacting sites to specific genomic elements such as promoter regions, exons or introns.

### Telomere Interactions Are Highly Enriched at Repetitive Genomic Elements

In our examination of Telomere-C peaks at previously identified TPE-OLD sites, we observed significant Telomere-C signals at nearby repetitive interstitial telomeric sequence (ITS) elements. This led us to hypothesize that telomeres might specifically interact with other repetitive elements across the entire genome. To test this, we analyzed Telomere-C peak overlaps with 5,590,282 unique repeats spanning 15,528 known RepeatMasker families (see Methods). Using shuffled peaks as a background control, we computed enrichment fold-change and significance (Fisher’s exact test and FDR). We identified several repeat families significantly associated with Telomere-C peaks (Figure 3A, Table 3, and Supplementary Table 1). This analysis revealed a broad and novel enrichment of telomeric interactions at repetitive elements. A notable hit included a family of telomeric tandem repeats (e.g., CTAACC and AGGGTT), which we collectively grouped as ITS elements due to their non-telomeric location. Unexpectedly, we also observed enrichment of Telomere-C peaks in Telomere-associated repeat 1 (TAR1) elements, which are often (but not exclusively) found in subtelomeres but do not contain any telomeric sequence. To our surprise, we also observed notable telomeric chromatin interactions with the relatively unknown repetitive element D20S16. While statistically significant ITS interactions were observed in every cell line tested except for IMR90 and WI38 cells, TAR1 interactions were found to be statistically significant in all cell lines (Figure 3B), establishing TAR1 as a universally interacting element.

**Figure 3.**
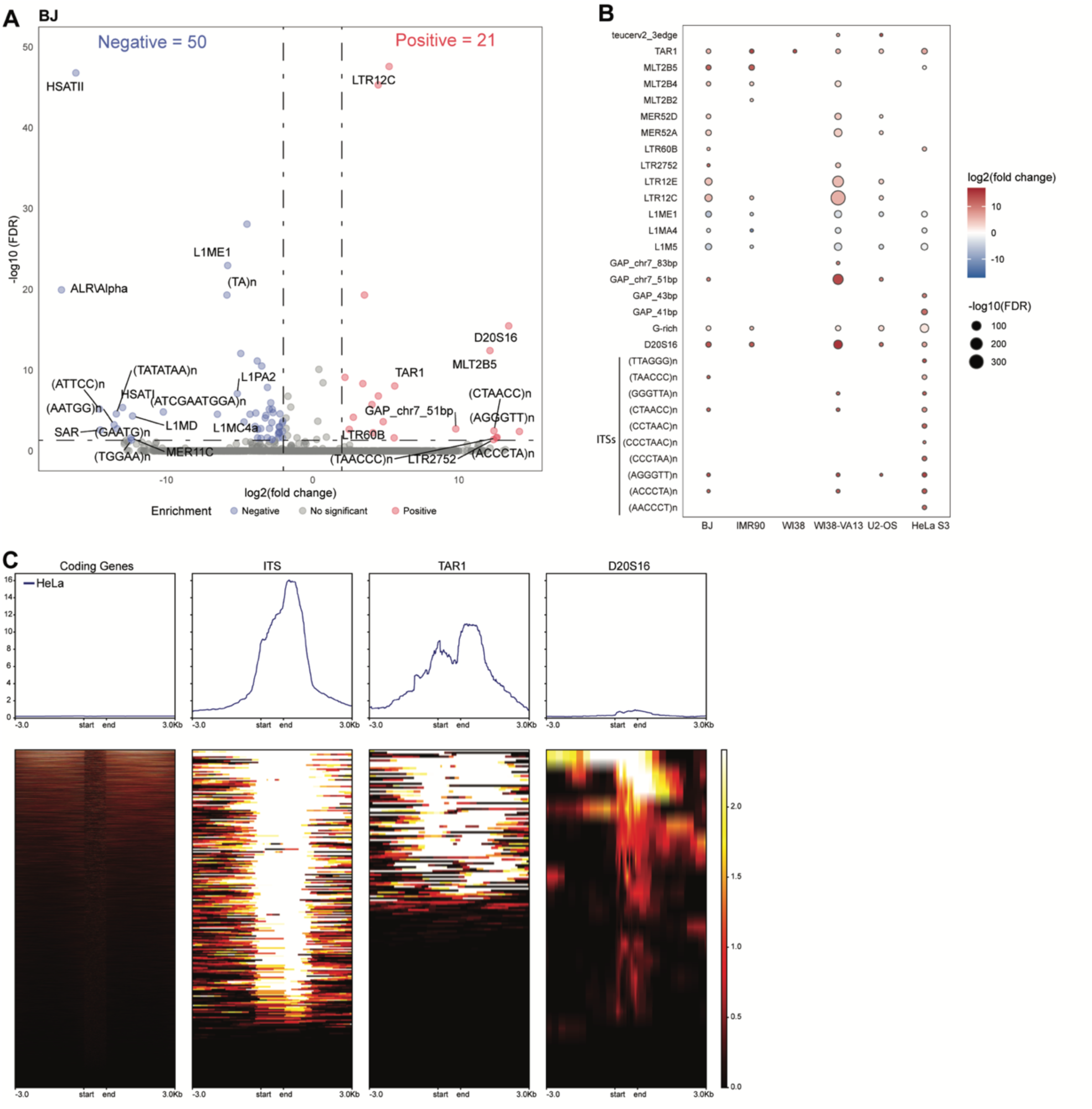
Enrichment of telomeric chromatin interactions at repetitive elements. **A** Volcano plot illustrating the fold enrichment of Telomere-C peaks (log_2_-scaled) versus the corresponding FDR (-log_10_-scaled) in BJ cells. Significantly enriched repeat families (FDR < 0.05 and fold change ≥ 4) are colored as red dots (positive enrichment) or blue dots (negative enrichment). **B** Bubble plot showing the top 30 significantly enriched repeat families across six cell lines. The color of the bubble corresponds to the fold change and the size of the bubble corresponds to the FDR. Blank cells indicate no significant enrichment in the corresponding cell (p < 0.05 by Fisher’s exact test). **C** Regional enrichment profile (top) and heatmaps (bottom) visualizing the enrichment of Telomere-C reads across regions of protein-coding genes, ITS, TAR1, and D20S16 elements, including 3 Kb of upstream and downstream flanking sequence, in HeLa S3 cells. The y-axis in the line plot represents the normalized read counts, while the heatmaps use a color gradient to indicate normalized read intensity.

**Table 3.**
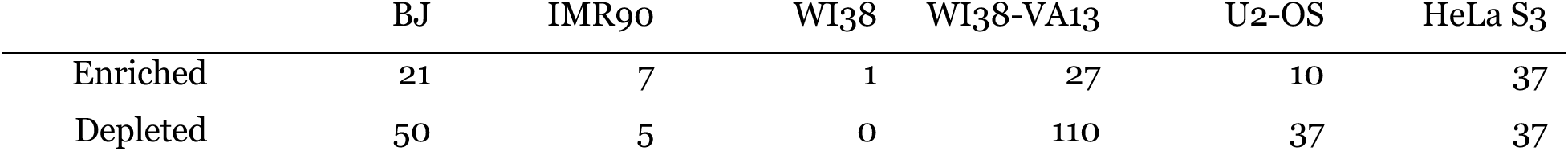
Summary of repeat family enrichment.

To investigate the regional distribution of telomeric interaction signal, we analyzed Telomere-C signals within each element and a 3 Kb flanking region, using HeLa S3 cells as a representative example (Figure 3C). The patterns of telomeric interactions varied dramatically based on the repeat type. For ITS elements, the signal covered the entire element and peaked in the downstream region. In contrast, TAR1 elements exhibited a bimodal signal, peaking in both upstream and downstream regions. This suggested that ITS and TAR1 repeats are organized via distinct telomeric chromatin interactions.

Overall, Telomere-C signals in protein-coding genes were significantly weaker than those affecting repetitive elements. This suggests that the ultra-long-range telomeric chromatin interactome predominantly engages genomic repeats, whereas previously reported interactions with protein-coding genes may be confined to specific cellular conditions and peri-telomeric sites as described for TPE and TPE-OLD. Finally, visualization of individual repeat elements across all cell lines revealed interesting cell-specific interaction patterns (Supplementary Figure 4). These findings fundamentally shift the understanding of telomeric 3D organization by demonstrating that genomic repeats, and not just gene promoters, constitute the major partners in the long-range telomere interactome.

### Telomere-ITS Interactions Exhibit Telomere-Maintenance-Dependent Clustering and Long-Range Contacts

To identify common and unique features of telomere-ITS interactions across the tested cell lines, we applied hierarchical clustering. This analysis revealed three distinct clusters of cell lines (Figure 4A and Supplementary 4A). The first cluster was characterized by diploid primary cells (BJ, IMR90, and WI38), the second was defined by ALT-positive cells (U2-OS and WI38-VA13) and the third cluster contained a telomerase-positive cell line (HeLa S3). This result indicated that ITS-telomere interactions are strongly associated with an underlying telomere maintenance mechanism.

**Figure 4.**
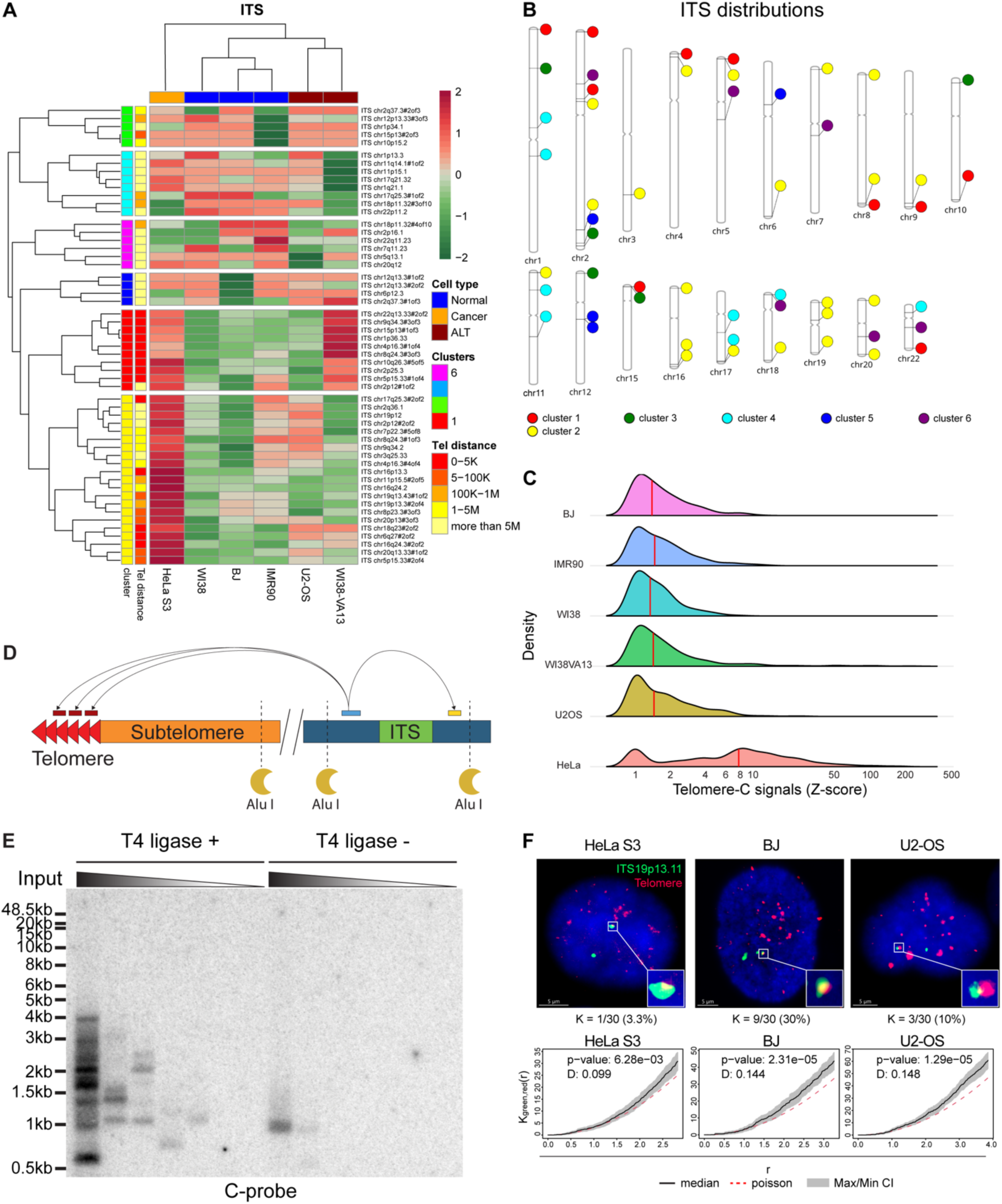
Validation of Telomere-ITS interactions. **A** Heatmap of binned raw Telomere-C signals. The intensity is color-coded from green (low, -2) to red (high, 2). ITS elements (rows) and cell lines (columns) are hierarchically clustered by Euclidean distance. Two colored sidebars on the left indicate the assigned ITS cluster (1-6, color-coded: red, yellow, green, cyan, blue, and purple). The other indicates distance to the telomere. A colored-bar at the top indicates cell types: normal (blue), cancer (yellow), and ALT+ (dark red). Only the top 10 most enriched ITS elements per cluster are displayed. **B** PhenoGram showing the distribution of ITS elements across the whole genome, color-coded by their assigned cluster as defined in **A**. **C** Ridgeline plot showing the density distribution of Telomere-C signal intensity in ITS regions across six indicated cells (the y-axis). The median is indicated in the red vertical bar. **D** Schematic of primer design to amplify proximal (ITS region) or distal (telomere-ITS) interactions (top). The red, blue, and yellow bars indicate the expected primer binding sites in the telomere, upstream of the ITS locus, and downstream of the ITS locus, respectively. The dashed lines represent the *AluI* restriction sites. **E** Visualization of Telomere-ITS18 amplicon by in-gel hybridization. The DNA template input was serially diluted (indicated above the gel) from 100 to 0.001 ng. The results were visualized using a ^32^P-labeled C-rich telomeric probe. **F** *Top:* 3D-FISH with scale bars (5 µm) for demonstrating colocalization between ITS19 (chr19q13.11, green) and telomeres (red) in HeLa S3, BJ and U2-OS. Nuclei were counterstained with DAPI (blue). The enlarged window represents a 5X magnification of selected colocalization foci. The number and percentage of colocalized nuclei are indicated below. *Bottom:* Pooled Ripley’s K-function plot representing aggregation levels between ITS19 and telomeres in HeLa S3, BJ, and U2-OS cells. The solid back line indicates the median of the bivariate K-function, with the grey shaded area indicating the confidence intervals. The red dotted line indicates the theoretical Poisson distribution (null hypothesis). The p-value and D statistics were calculated using Kolmogorov– Smirnov test

Clustering of individual ITS elements demonstrated that the interaction distance exhibits cell type specificity (Figure 4A-B). Specifically, telomere-ITS interactions preferentially occurred near the ends of chromosomes in cancer cell lines (cluster 1 in HeLa S3, U2-OS, WI38-VA13, and cluster 2 in HeLa S3), whereas in normal cells, they were more commonly found at ultra-long-range distances (> 5 Mb) from the telomere. This differential interaction distance is notable, given that >80% of overall telomeric interactions occur at the ultra-long-range. We also observed unusually strong telomere-ITS interactions in HeLa S3 cells compared to the other cell lines, suggesting a potential link between telomere shorting, telomerase reactivation and the stability of telomere-ITS interactions (Figure 4C).

To validate the existence of these long-range telomeres-ITS contacts, we performed traditional Chromatin Conformation Capture (3C) followed by PCR and Southern blotting. We selected two ITS loci in BJ cells, hereafter referred to as ITS18 (on chromosome 18) and ITS19 (on chromosome 19), for which primer pairs could be designed within the same AluI restriction fragment (Figure 4D). ITS primers were paired with the telomeric tel1 primer, which is a widely used primer to amplify telomeric repeats without primer dimer-derived products^42^. Since tel1 binds to all possible 5-repeat telomeric hexamers, the expected PCR products vary in size.

We observed a laddered smear of PCR products using both the tel1-ITS18 and tel1-ITS19 primers in 3C DNA with proximity ligation (Supplementary Figure 5A). In contrast, no smear was detected in 3C DNA without proximity ligation or in a non-template control experiment, verifying that the PCR products are specific to the long-range chromatin interaction (Supplementary Figure 5B). To exclude the possibility of PCR amplification bias, we perform ^32^P-labeled telomeric C-probe in-gel hybridization following a dilution strategy similar to TeSLA (see Methods)^43^. Confirming our Telomere-C findings, we observed multiple discrete bands only in proximity ligated 3C DNA (Figure 4E and Supplementary Figure 5C). These results collectively confirm the existence of a long-range interaction between the ITS18/19 elements and telomeres.

Finally, we used 3D-Fluoresence In Situ Hybridization (FISH) to provide further validation for the remote interactions between telomeres and ITS elements, combining probes targeting ITS19 and telomeric DNA (Figure 4F). Analyzing at least 30 nuclei per cell line, we detected colocalization events in *n* = 1 (3.3%), 9 (30%), and 3 (10%) nuclei in HeLa S3, BJ, and U2-OS cells, respectively.

To rigorously determine if the observed ITS and telomere foci were spatially associated, we applied Ripley’s K-function. Our statistical analysis showed that all cell lines exhibited aggregation of ITS19 and telomeric foci exceeding a theoretical Poisson distribution (p-value < 0.05). Compared to BJ and U2-OS, HeLa S3 cells showed a lower statistical power (p-value: 6.28x10-3), smaller D statistic (0.099), and fewer direct colocalization events (3.3%). This result is consistent with our Telomere-C finding that HeLa S3 cells exhibit a lower preference for telomere-ITS interaction at the ultra-long-range, given that ITS19 is located over 5 Mb away from the chromosome 19 telomere.

### Telomere-TAR1 Interactions Show Cell Lineage-Specific Clustering and Stable Peri-telomeric Enrichment

We next extended our analysis to TAR1 repeat elements, which, unlike ITS elements, do not contain TTAGGG or degenerate telomeric sequences but were consistently and significantly enriched with Telomere-C peaks across all the cell lines tested (Figure 3B). Performing hierarchical clustering of Telomere-C signal in individual TAR1 elements showed that the enrichment of TAR1-telomere interactions varied by cell origin (Figure 5A and Supplementary Figure 4B). For example, the ALT-positive cell line WI38-VA13 and their non-immortalized primary diploid parental cell, WI38, formed one cluster, while the other primary fibroblast cell lines BJ and IMR90 formed a separate cluster. This pattern suggests that TAR1-telomere organization is associated with cell lineage or origin rather than solely the telomere maintenance mechanism.

**Figure 5.**
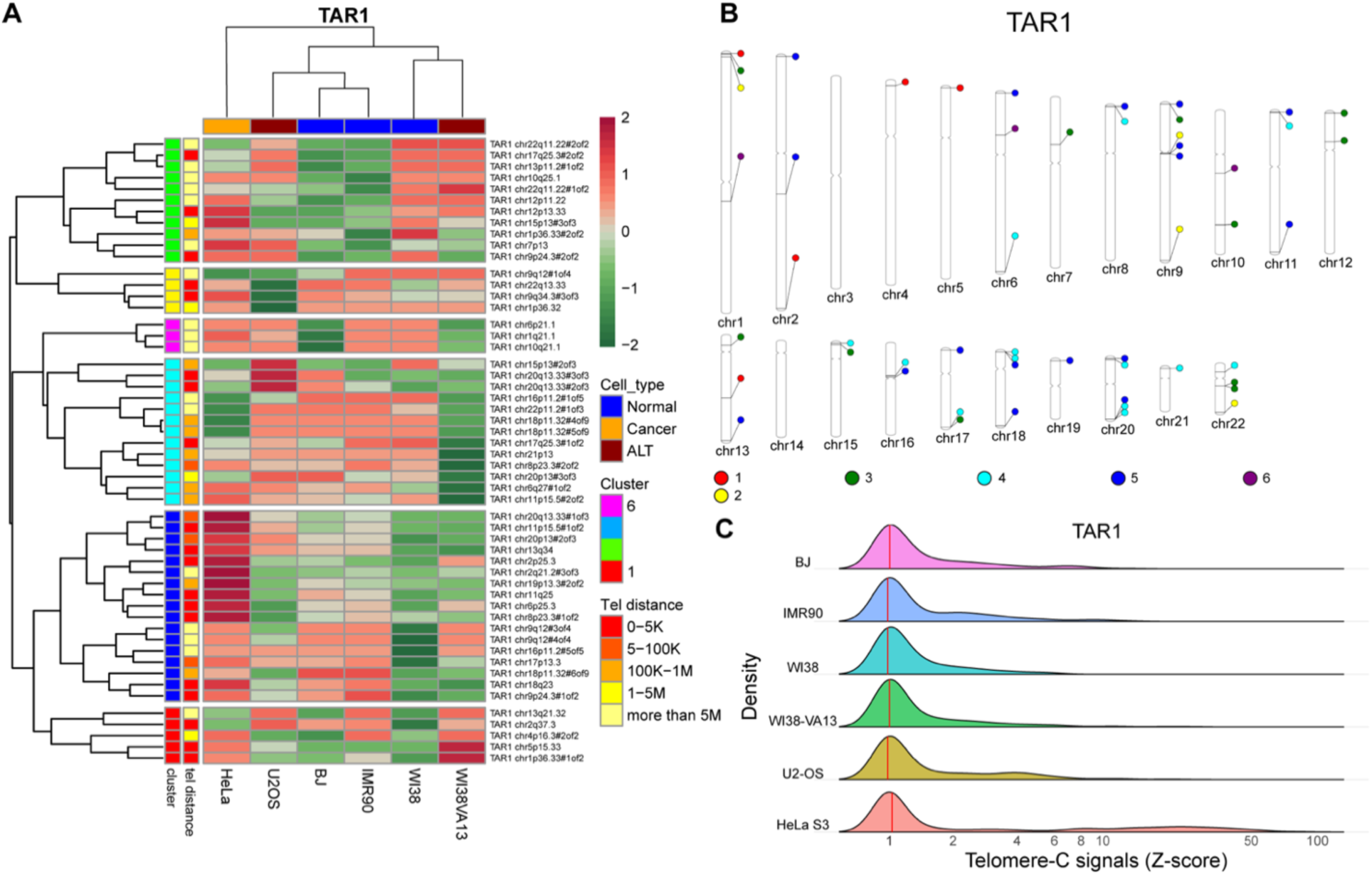
Telomere-TAR1 interactions are preferentially adjacent to telomere-rich regions. **A** Heatmap of normalized Telomere-C signal intensity (Z-score) is color-coded from green (low, -2) to red (high, 2). TAR1 elements (rows) and cell lines (columns) are hierarchically clustered by Euclidean distance. Two colored sidebars on the left indicate the assigned TAR1 cluster (1-6, color-coded: red, yellow, green, cyan, blue, and purple) and the distance to the telomere. A colored bar at the top indicates cell types: normal (blue), cancer (yellow), and ALT+ (dark red). Only the top 10 most enriched TAR1 elements per cluster are displayed **B** PhenoGram showing the distribution of TAR1 elements, color-coded by their assigned cluster as defined in **A**. **C** Ridgeline plot showing the density distribution of Telomere-C signal intensity in TAR1 regions across six indicated cells (the y-axis). The median is indicated in the red vertical bar.

In contrast to the findings for the overall interactome, we observed that most of the strong Telomere-C signals occurred at TAR1 elements located within 1 Mb of chromosomal ends, which reflects the known peri-telomeric distribution of TAR1 elements in the human genome (Figure 5B and Supplementary Figures 6 A-B). Additionally, and unlike ITS interactions, we did not observe a particular enrichment of interaction strength in HeLa S3 telomerase-positive cells compared to other cell types (Figure 5C). This suggests that the TAR1-telomere interactions represent a stably maintained, peri-telomeric organization common across diverse cell types.

### D20S16-Telomere Interactions Are a Distinctive Feature of the ALT Phenotype

In addition to ITS and TAR1, our unsupervised repetitive element analysis also revealed a strong and significant enrichment of Telomere-C signal in D20S16 elements, with *n* = 5 tested cell lines exhibiting a higher fold-enrichment compared to other repetitive elements (Figure 3B). D20S16 was initially discovered as polymorphic satellite DNA on chromosome 20, though its function remains poorly understood^44^. Recent studies found that D20S16 exhibited high expression levels during early development and in breast cancer cells^45–47^.

Clustering analysis demonstrated that D20S16-telomere interactions exhibited an even stronger cell-type specificity than ITS and TAR1 interactions (Figure 6A and Supplementary Figure 4C). ALT-positive cells (WI38-VA13 and U2-OS) formed a separate group and showed the strongest enrichment by far, whereas diploid normal cells (BJ, IMR90, and WI38) and hTERT-positive cancer cells (HeLa S3) each formed their own distinct clusters. This finding is particularly notable because D20S16 elements exist in large clustered arrays on eight chromosomes and are positioned far away from the telomere, contrasting sharply with the smaller, sometimes peri-telomeric, ITS and TAR1 elements (Figure 6B and Supplementary Figures 6C and 7A).

**Figure 6.**
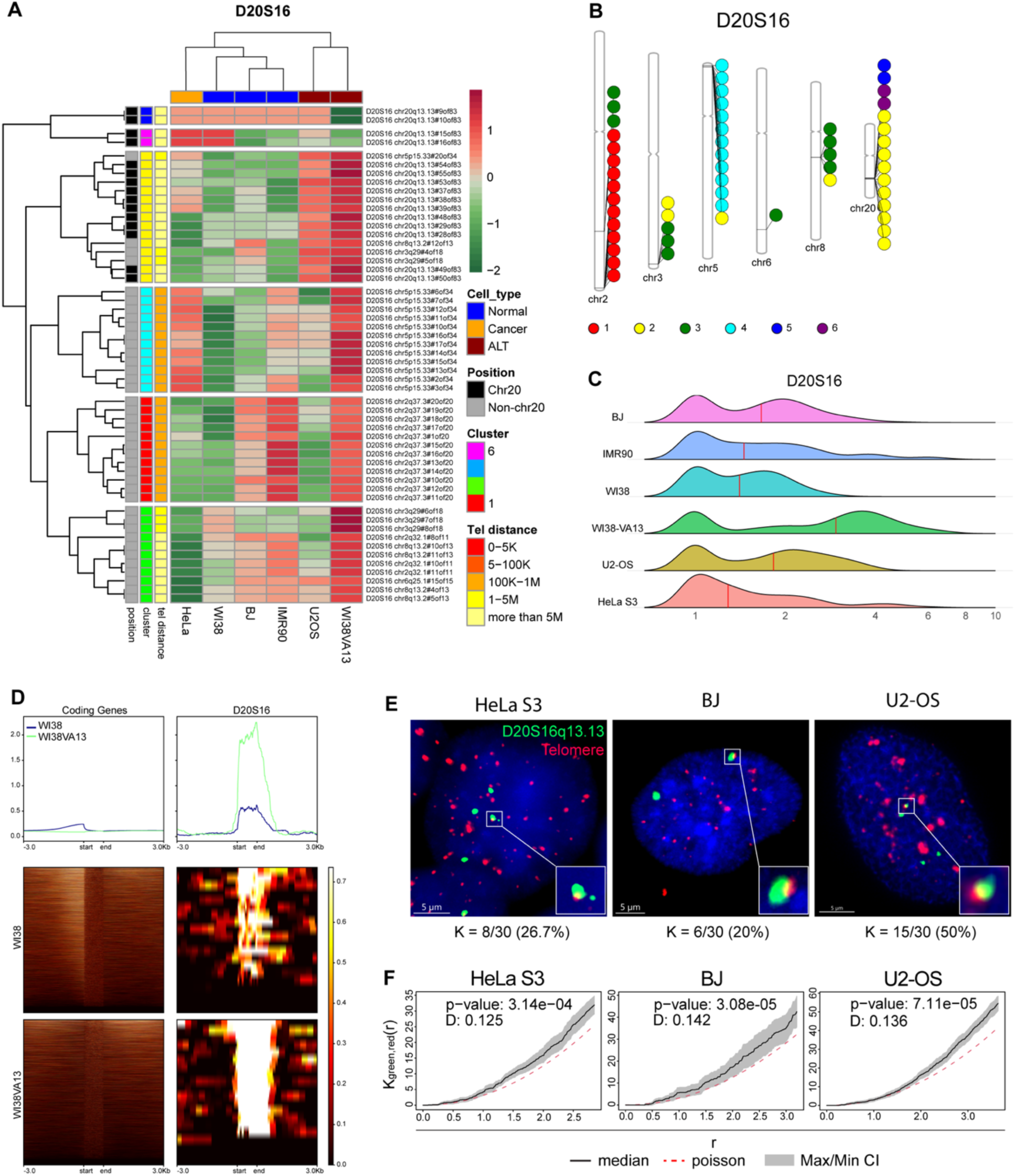
Telomere-D20S16 interaction is positively associated with the ALT phenotype. **A** Heatmap of normalized Telomere-C signal intensity (Z-score) color-coded from green (low, -2) to red (high, 2). D20S16 elements (rows) and cell lines (columns) are hierarchically clustered by Euclidean distance. Three colored sidebars on the left indicate the assigned D20S16 cluster (1-6, color-coded: red, yellow, green, cyan, blue, and purple), distance to the telomere and chromosome. A colored-bar at the top indicates cell type: normal (blue), cancer (yellow), and ALT+ (dark red). Only the top 10 most enriched D20S16 elements per cluster are displayed **B** PhenoGram showing the distribution of D20S16 elements color-coded by their assigned cluster as defined in **A**. **C** Ridgeline plot showing the density distribution of Telomere-C signal intensity in D20S16 regions across six indicated cells. The median is indicated in the red vertical bar. **D** Read enrichment profile (top) and heatmaps (bottom) visualize the enrichment of Telomere-C reads in D20S16 elements, including a 3 Kb upstream and downstream flanking region in WI38 and WI38-VA13 cells. The y-axis in the line plot represents the normalized read counts, while the heatmaps use a color gradient to indicate normalized read intensity. **E** 3D FISH images with scale bars (5 µm) demonstrating colocalization between D20S16 (chr20q13.13, green) and telomeres (red) in HeLa S3, BJ, and U2-OS cells. Nuclei were counterstained with DAPI (blue). The enlarged window represents a 5X magnification of selected colocalization foci. The number and percentage of colocalized nuclei are indicated below. **F** Pooled Ripley’s K-function plot illustrating the aggregation levels between D20S16 and telomeres in HeLa S3, BJ, and U2-OS cells. The solid back line indicates the median of the bivariate K-function, with the grey shaded area indicating the confidence intervals. The red dotted line indicates the theoretical Poisson distribution (null hypothesis). The p-value and D statistics were calculated using a Kolmogorov– Smirnov test.

Importantly, we observed a significantly higher Telomere-C intensity in the ALT-positive WI38-VA13 and U2-OS cell lines compared to other lines tested, strongly suggesting that D20S16-telomere interactions are an important, distinctive feature of the ALT phenotype (Figure 6C). To definitively ascertain that the telomere-D20S16 interaction is ALT-specific, we compared the ALT-positive WI38-VA13 cell line directly to its diploid parental counterpart, WI38 (Figure 6D). This analysis clearly demonstrated that WI38-VA13 cells exhibited a stronger D20S16 Telomere-C signal compared to WI38, a signal that significantly outweighed the average telomeric interactions with protein-coding genes.

To validate the newly discovered ALT-associated telomere-D20S16 interaction, we performed FISH by co-hybridizing telomere and D20S16-q13.13 (or D20S16-q13.13-2) probes, targeting two clusters of D20S16 elements on chromosome 20 that exhibited strong Telomere-C signal in ALT-positive cells. Analyzing 30 nuclei per cell line, we found that the frequency of co-localization events correlated with the ALT phenotype in the larger primary D20S16 cluster (D20S16-q13.13): *n* = 15 (50%) of U2-OS nuclei exhibited co-localization, compared to *n* = 8 (26.7%) in HeLa S3 and *n* = 6 (20%) in BJ cells (Figure 6E and Table 4). A second, smaller, cluster of D20S16 elements (D20S16-q13.13-2) showed significant co-localization with telomeres, although this did not appear to be particularly related to ALT status (Supplementary Figure 7B and Table 4).

**Table 4.**
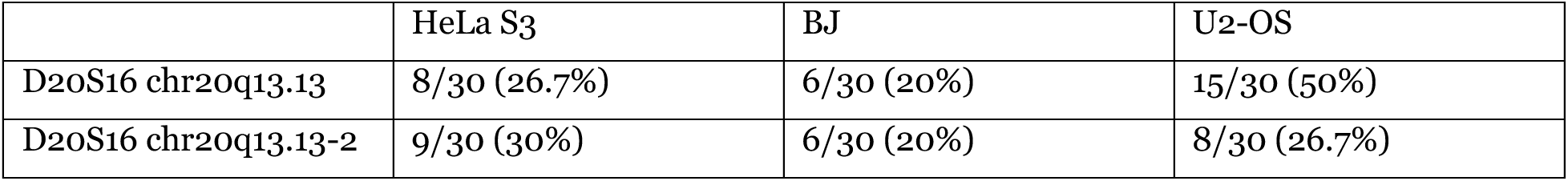
Summary of Telomere-D20S16 colocalization events by FISH.

We next applied Ripley’s K-function and a Kolmogorov–Smirnov test to evaluate spatial aggregation. Our analysis showed that all tested cell lines exhibited aggregation of D20S16 elements and telomeres (Figure 6F and Supplementary Figure 7C). Notably, while all cell lines passed the statistical threshold, BJ cells demonstrated relatively lower statistical power and smaller effect size, consistent with the lower frequency of co-localization events in BJ (20%) compared to HeLa S3 and U2-OS (26.7% and 50%, respectively). These results establish the D20S16-telomere interaction as a specific and robust ALT biomarker.

In summary, we developed Telomere-C to construct high-resolution, genome-wide maps of telomeric chromatin interactions in human cells, providing a definitive solution to a long-standing technical challenge. Using our approach, we were able to recapitulate and precisely pinpoint the location of known telomere-gene interactions involved in TPE. Unexpectedly, we discovered that telomeres physically interact with remote regions over 5 Mb away from chromosome ends, and that this ultra-long-range interactome is strongly enriched at repetitive elements, in particular ITS, TAR1, and D20S16 elements. The preference of each repeat interaction varied significantly by cell type and telomere maintenance mechanism, suggesting that long-range telomere-repeat interactions are of profound phenotypic relevance. Notably, we discovered a strong enrichment of D20S16 interactions specifically in ALT-positive cancer cell lines, establishing it as a novel, robust biomarker and potentially mechanistic driver for this pathway. Overall, our study offers an important foundation for future studies of telomeric chromatin interactions. These findings fundamentally expand our understanding of telomeres from mere end protection structures to central components of genome-wide chromatin organization, which we believe will ultimately prove critical in resolving complex structural variation involving telomeres in diseases such as cancer.

## Discussion

We developed Telomere-C, a novel capture-based approach to measure telomeric 3D chromatin organization. Our approach achieves a high sensitivity for telomeric chromatin interactions by enriching for telomere-associated DNA using a biotinylated probe while depleting the uninformative telomeric sequences themselves post-capture. This ligation-free method drastically reduces mapping complexity, enabling high-resolution studies of repetitive genomic regions. Telomere-C exhibited significantly enhanced efficiency and resolution compared to Hi-C and Micro-C. In IMR90 cells, it yielded millions of telomere-associated reads from ∼40 million total reads, vastly outperforming the few thousand reads recovered by conventional methods. Using this robust platform, we constructed the first high-resolution maps of the genome-wide telomeric interactome across normal, telomerase-positive, and ALT-positive cell lines.

With Telomere-C, we discovered a new class of interaction that extends far beyond previously defined long-distance interactions, which we have termed ultra-long-range telomeric chromatin interactions (>5 Mb). Notably, these were the most abundant type of interaction in most cell lines, often constituting more than 80% of all interactions. These interacting loci were largely cell type specific, suggesting a unique spatial organization landscape in each cell type. We observed that the proportion of ultra-long-range interactions in IMR90 (43%) and WI38 (45%) was lower than in other cell lines. This difference likely reflects a change in telomeric organization driven by cell status (e.g., transformation or maintenance mechanism) and warrants further investigation.

Our high-resolution data provided physical evidence supporting previous studies that proposed the telomere position effect can regulate gene expression at long distances (TPE-OLD)^20,22,25^. We not only validated known telomeric interactions at the *TERT*, *ISG15*, and *DUX4* loci but were able to pinpoint these interactions to specific genomic elements such as exons, introns, and proximal regions of transcripts. While these findings physically support the TPE-OLD model, this study is limited by the absence of transcriptional data in cells with variable telomere lengths or perturbed binding sites. Therefore, whether these interactions at these loci directly influence gene expression remains an uncovered question for future work

The enrichment of Telomere-C signal at ITS and TAR1 repeat elements in subtelomeric and intra-chromosomal regions was markedly stronger than at protein-coding genes, fundamentally expanding the known partners of the telomeric interactome. The mechanism of these interactions remains elusive. Previous studies suggest shelterin components like *TRF2* could mediate telomere-ITS interactions, as TRF2 has been shown to bind ITS elements^2,48,49^. Alternatively, the telomeric long non-coding RNA *TERRA* may be involved by recruiting HP1α and increasing the H3K9me3 density or by forming DNA-RNA hybrid R-loops at telomeres^50,51^. While it is attractive to speculate that *TERRA* might also form R-loops at ITS elements, this is currently unvalidated. The co-localization we observed between ITS and telomeres, combined with prior reports of ITS elements contacting nuclear lamina components, suggest telomere-repeat interactions have broad implications in both cis and trans nuclear organization^52^. Future mechanistic studies are necessary to define the roles of TRF2 and TERRA in mediating these widespread ITS and TAR1 interactions.

Most notably, we observed specific telomeric chromatin interactions with the D20S16 satellite on chromosome 20 that are enriched in ALT-positive cell lines, suggesting a novel mechanism for ALT maintenance. D20S16’s expression pattern (high in early development and cancer) is similar to that of key oncogenes, suggesting a contribution to oncogenesis^46,47^.

Based on our results and existing literature, we hypothesize that the Telomere-D20S16 interaction is mediated via orphan nuclear receptors (NRs), which are known to serve as a molecular hub for ALT recombination by bridging ALT telomeres to distal genomic regions^30,34^. This interaction is proposed to drive targeted telomere insertion, a key feature of the ALT-associated genomic instability^33^. Further investigation is needed to validate this proposed mechanism, including proteomic analysis to identify binding partners of D20S16 and sequencing the interacting sites to confirm the presence of the ALT-specific sequences.

In closing, we analyzed genome-wide telomeric chromatin interactions in various human cell lines and discovered a new class of ultra-long-range telomeric chromatin interactions, which are the most abundant type of interaction in the cell lines investigated. Furthermore, we expanded our understanding of telomeric chromatin interaction by showing that the major interacting elements are not the promoters of protein-coding genes but rather repetitive elements, most notably D20S16 satellites in ALT-positive cancer cells. The identification of this extensive telomeric interactome suggests that telomeres are integral hubs for higher-order chromatin folding. This functional expansion provides a necessary framework for investigating their involvement in driving complex telomere-associated structural variation in cancer.

## Material and Methods

### Cell culture

BJ, U2-OS, HeLa S3 and IMR-90 cells were gifts from Dr. Michael Berens and Dr. Haiyong Han at the Translation Genomic Research Institute. These cell lines, along with WI38 and WI38-VA13, were originally obtained from American Type Culture Collection (ATCC). BJ and IMR90 were authenticated by STR profiling at ATCC. All the cell lines, except U2-OS, were cultured in Eagle’s Minimum Essential Medium (ATCC, Cat. No. 30-2003) while U2-OS cells were cultured in McCoy’s 5A medium (ATCC, Cat. No. 30-2007). Both mediums were supplemented with 10% FBS (VWR, Cat. No. 97068-091) and cell lines were cultured at 37°C with 5% CO2 in humidified incubators, according to ATCC subculturing procedures.

### Cell harvesting and dual crosslinking

Cells were harvested using 0.25% Trypsin and pelleted by centrifugation at 2,500 rpm. At least 6 million cells were crosslinked with 1.5 mM ethylene glycol bis(succinimidyl succinate) (Thermo Scientific, 21565) for 30 minutes followed by adding 1% formaldehyde (Thermo Scientific™ Pierce, PI28906) for 10 minutes. The crosslinking reaction was quenched by the addition of 200 mM glycine. Crosslinked cells were then snap frozen into individual pellets (containing 3 million cells each) and stored at -80°C in 1.5 mL tubes.

### Telomere Conformation Capture by sequencing (Telomere-C)

Frozen crosslinked cell pellets were thawed on ice and resuspended with cell lysis buffer (50 mM HEPES, pH 7.5, 150 mM NaCl, 1 mM EDTA, 1% Triton X-100 (wt/vol), 0.1% sodium deoxycholate (wt/vol)) containing 0.1% SDS (wt/vol) and 1X protease inhibitor (Roche, 4693116001), then placed on a tube revolver (Fisherbrand, 88881051) at 20 rpm for 20 minutes at 4°C. The mixture was refreshed with cell lysis buffer and incubated for another 10 minutes at 4°C. Lysed pellets were incubated with nuclear lysis buffer (cell lysis buffer containing 1% (wt/vol) SDS and 1X protease inhibitor) for 15 minutes at 4°C, twice. The lysed pellets were resuspended in Qiagen EB buffer (Qiagen, 19086), transferred to a microTUBE (Covaris PN 520045), and sonicated using a Covaris E220 Focused-Ultrasonicator with the following parameter settings: 105W Peak Incident Power, 2% Duty Factor, 200 Cycles per burst, 240 second duration time, and temperature limit from 4-9°C . The fragmented chromatin was centrifuged at full speed for 10 minutes at 4°C and then supernatant was transferred into a new 1.5 mL tube. For quality control, a 15 µL of supernatant was purified using the Zymo ChIP DNA Clean kit (Zymo PN: D5205), then subjected to Qubit 4 Fluorometer (Invitrogen, Q33238) and 4150 TapeStation (Agilent, G2992AA) for assessing quantity and sizes (peaked at 1000 bp).

Fragmented chromatin was concentrated with an Amicon Ultra-0.5mL centrifugal filter device, 100 kDa (Millipore, UFC510008). The concentrated chromatin was subjected to a pre-cleaning step by incubating it with Dynabeads MyOne Streptavidin T1 (Invitrogen, 65601) in LBJDLS buffer (10 mM HEPES pH 7.5, 100 mM NaCl, 2 mM EDTA, 1 mM EGTA, and 0.2% (wt/vol) SDS) containing 1X protease inhibitor. The mixture was rotated for 2 hours at 4°C. Following incubation, the tube was placed on a magnet stand, and the pre-cleaned chromatin was transferred to a clean 1.5 mL microtube. A 15 µL aliquot of supernatant was taken, purified, and measured for DNA concentration by Qubit 4. The remaining sample was stored as the input control for subsequent Illumina library preparation.

The pre-cleaned chromatin was mixed with 250 nM biotinylated PNA probe (CCCTAA repeats; PNA Bio, F2001) and incubated using a thermocycler with the following program: 25 °C for 3 minutes, 71 °C for 9 minutes, 38 °C for 1 hour, 25°C to cooling down. The hybridized complex was captured by incubating Dynabeads MyOne Streptavidin T1 in LBJD buffer (modified LBJDLS buffer with 30 mM NaCl) for 90 minutes at room temperature. The beads were then washed three times with LBJDLS buffer. The bead-bound chromatin was resuspended in TE buffer, followed by proteinase K treatment for 4 hours at 65°C. The supernatant (containing telomere-associated DNA) was recovered after placing the mixture on a magnetic stand and transferred to clean 2 mL microtubes. The recovered DNA was purified using phenol-chloroform extraction.

The purified DNA and the input control DNA samples were digested with AluI and indexed with the IDT for Illumina TruSeq UDI-UMI adaptors using the KAPA HyperPrep Kit (Roche, 7962363001), according to the manufacturer’s instructions. The amplified libraries were size-selected using AMPure XP beads (Beckman Coulter, A63881) via a double size selection at 0.8X and 0.56X beads ratios, respectively. Library quality control was performed using the 4150 TapeStation for size distribution (peaked at 500 bp) and the Qubit 4 Fluorometer for concentration. The final libraries were sequenced on an Illumina NovaSeq X instrument.

### Data processing of Telomere-C data

Sequencing data were processed using our Telomere-C Snakemake pipeline, available on our GitHub (https://github.com/barthel-lab/Telomere-C). Briefly, UMI barcodes were first extracted using UMI-tools (v1.1.1). The reads were then subjected to adapter trimming using Cutadapt (v4.8)^53,54^. Illumina adapters were marked using GATK4 (v4.2.2.0), and the processed reads were subsequently mapped to the human reference genome CHM13v2 using BWA-MEM (v0.7.17)^55–57^. Following alignment, UMI deduplication was performed using UMI-tools. The reads were filtered using BEDtools (v2.30.0) and SAMtools (v1.13)^58–60^. Specifically, all reads mapping to the exclusion regions (T2T.excluderanges) were discarded, with the exception of reads aligning to the telomeric regions defined in chm13v2.0_telomere.bed (T2T consortium).

### Telomere-C peak calling and visualization

Probe captured reads were normalized for GC content, sequencing depth, and input control data using an in-house Python script. Peak calling was performed on the normalized data using the RGT (v1.0.0)^61^. The normalized data and resulting called peaks were visualized using karyoploteR (v1.28.0)^62^ or the IGV^63^.

### Analysis of Telomere-C peaks

The size distribution of peaks was plotted as a box plot using the R package ggplot2 (v3.5.2). The distance between Telomere-C peaks and the nearest telomeres (or centromeres) was computed, based on the annotations from the T2T Consortium. These distances were visualized in a density plot, or converted to proportions for display in a pie chart. To analyze the uniqueness and overlap of Telomere-C peaks, the intersections of peak regions among all cell lines were determined using the *merge* function from BEDtools. The results were visualized in an UpSet plot using the R package, UpSetR (v1.4.0)^64^.

### Hi-C

Frozen crosslinked cell pellets were used for Hi-C library preparation using the Arima High Coverage HiC Kit (Arima Genomics, Cat. No. A101030) and Arima Library Prep Kit v2 (Arima Genomics, Cat. No. A303011), according to the manufacturer’s instructions. 1.5 µg of proximity-ligated DNA suspended in 130 µL of the elution buffer was added to a microTUBE (Covaris, Cat. No. 520045) and sonicated using a Covaris E220 with the following parameters: Temperature (4-7 °C), Peak Incident Power (105W), Duty Factor (5%), Cycles per Burst (200), Treatment time (75 s). Post-sonication QC was performed on a TapeStation with HSD5000 screentape (Agilent Technologies, Cat. No. 5067-5592) to verify that fragment sizes were between 550-600 bp prior to size selection. Size selection was performed with AMPure XP beads (Beckman Coulter, Cat. No. A63881) to collect fragments >400 bp. 200 ng of size-selected DNA was subjected to biotin enrichment. The enriched products were end-repaired, adapter-ligated, and tagged with manufacturer supplied P7 and P5 8 bp index pairs using 5 cycles of PCR. The final bead-purified libraries were quantified by Qubit dsDNA HS (Thermo Fisher, Cat. No. Q33231) and TapeStation with HSD5000 screentape, pooled, and re-quantified. The pool was sequenced on a NovaSeqX instrument using a 10B flow cell (Illumina, Cat. No. 20085594) for 151 x 151 cycles with a 1% PhiX spike-in, aiming for generating 25x coverage.

### Data processing of Hi-C and Micro-C

Micro-C data was obtained from the NCBI Gene Expression Omnibus (GEO) with accession number GSM7661134. Both Hi-C and Micro-C data were processed using our Snakemake pipeline (https://github.com/barthel-lab/micro-C_data_processing), which is adapted from the Dovetail Genomics pipeline (https://micro-c.readthedocs.io/en/latest/fastq_to_bam.html). Briefly, to analyze telomere-associated reads, reads containing at least one 60-mer of (TTAGGG)_10_ in either read 1 or read 2 were first retained using KMC (v3.2.4). The filtered reads were mapped to the human reference genome CHM13v2 using BWA-MEM (v0.7.17) with the *-5SP -T0* options. Ligation junctions were parsed using *pairtools parse* with *--walks-policy all* option (v1.1.2). Given the repetitive nature of telomere-associated reads, one end of a read pair is unmappable. *pairtools select* was performed using the filter *’(mapq1 >= 40) or (mapq2 >= 40)’* to retain read pairs where at least one read passed the mapping quality threshold. The filtered reads were then sorted and deduplicated using *pairtools dedup* with *--output-unmapped* option, which allowed for the retaintion of pairsam files containing unmappable reads. These files were then converted into BAM format using pairtools split with *--output-sam* option followed by sorting with *samtools sort.* The sorted BAM files were converted to bigwig format using *bamCoverage* from Deeptools (v3.5.4)^65^.

### Enrichment of Telomere-C peaks in repeat families

The analysis and visualization were performed by our in-house scripts. For each repeat family (based on RepeatMasker v4.1.2p1.2022Apr14 annotation), 𝑚 loci in the family were analyzed to compute the number of interaction events between Telomere-C peaks and the repeat loci were calculated in the same method. The total number of overlap bases was calculated as 𝐿 = ∑_1_^m^ 𝐿*_i_*. The total bases of the repeat family was denoted as 𝑀. To establish the background, the Telomere-C peaks were shuffled once using the *shuffle* function from BEDtools. The total number of overlap bases was calculated as 𝐿𝐵 = ∑_1_^m^ 𝐿*_Bi_*. Using these values 𝐿, 𝐿𝐵, and 𝑀, the enrichment of each family was calculated by log_2_(𝐿 / 𝐿𝐵,), and Fisher’s exact test was applied to calculate the p-value. The False discovery rate (FDR) was computed across the entire list of families. Results were visualized in a bubble plot to illustrate FDR and enrichment of repeat families across six cell lines. In addition, the analysis results for BJ Telomere-C were also visualized in a volcano plot as the representative results.

### Comparison of Telomere-C coverages in the repeat elements across cells

The coverages of normalized Telomere-C reads within the loci of ITS, TAR1, or D20S16 repeat elements were computed using the *multiBigwigSummary* function from Deeptools (v3.5.4). The density distribution of Telomere-C coverages was visualized in a ridgeline plot using the R package ggridges (v0.5.6).

To compare the coverages across six cell lines, the coverages (excluding zero read counts) were first transformed to a 𝑙𝑜𝑔_2_ scale and then converted to a Z-score. Specifically, for each locus, we compute the mean (𝜇) and standard deviation (𝜎) of the 𝑙𝑜𝑔_2_coverage across all cell lines, defining Z score as:

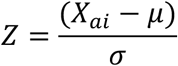

Here, 𝑋_ai_ indicates 𝑙𝑜𝑔_2_coverage at the 𝑎 locus in the 𝑖 cell. The data were visualized in a heatmap using the R package pheatmap (v1.0.13)^66^ with annotations. Specifically, we performed hierarchical clustering and manually defined six clusters by cutting the dendrogram using the *cutree* function. The heatmap was annotated with three tacks showing the distance to telomeres, cluster group, and position on chromosome 20.

To illustrate the genomic distribution of the analyzed repeat elements, the loci and corresponding cluster group were visualized using a PhenoGram Plot^67^.

### Chromatin conformation capture (3C)

The 3C was performed as described previously with modifications. Briefly, crosslinked cells were treated with cell lysis buffer and nuclear lysis buffer^68^. Lysed pellets were resuspended in 1X rCutSmart buffer (NEB, B6004S) containing 0.1% SDS and incubated for 10 minutes at 65°C followed by 10% Triton X-100 for quenching. AluI (520U, NEB, R0137L) was added, incubated overnight at 37°C, and then inactivated by incubating for 20 minutes at 65°C. The ligation mix, containing 1X ligation buffer (Invitrogen, 46300018), 50U T4 ligase (NEB, M0202L), 1% Triton X-100, 100 µg/ml BSA (Invitrogen, AM2614), was added to digested chromatin and incubated for 4 hours at 16°C. 60 µl of 10 mg/ml Proteinase K were added to the ligation product and incubated overnight at 65°C. The DNA purification was performed by phenol-chloroform extraction followed by ethanol precipitation and resuspended in Qiagen EB buffer. The 100 µg of purified 3C DNA was used as a template for PCR. Primers were designed to target AluI fragments containing ITS on chr18:15,145,858-15,146,709 and chr19:19,306,848-19,307,366 (CHM13v2) paired with tel1 primer, which targets telomeric hexamer repeats without primer dimer-derived products^42^. Specifically, tel1 paired with ITS19RC and tel1 paired with ITS18RC were used to investigate Telomere-ITS18 and Telomere-ITS19 distal interaction, respectively. ITS18-Fw paired with ITS18-Rev and ITS19-Fw paired with ITS19-Rev were used to amplify loci of ITS18 and ITS19 as positive control, respectively. The amplified products were visualized using electrophoresis on a 1% agarose gel containing 1X SYBR™ Safe DNA Gel Stain (Invitrogen, S33102) or by Southern blotting.

Primer sequences are described below:

tel1:

5’-GGTTTTTGAGGGTGAGGGTGAGGGTGAGGGTGAGGGT-3’

ITS18-Fw:

5’-CGTGACCCTGACCTTTACCCAT-3’

ITS18-Rev:

5’-TCCTAACGAGGTTCTCCCCA-3’

ITS18-RC:

5’-ATGGGTAAAGGTCAGGGTCACG-3’

ITS19-Fw:

5’-TTTGCATACAGGGCGTCACC-3’

ITS19-Rev:

5’-AGCCCCGTCTTGCAGTCTTT-3’

ITS19-RC:

5’-GGTGACGCCCTGTATGCAAA -3’

### Southern blotting

The PCR products were subjected to constant-field gel electrophoresis, which were separated using the Sub-Cell GT Horizontal Electrophoresis System (BioRad) with 0.7% agarose gel in 1× TAE buffer at room temperature. The gel was dried for 3hrs at room temperature. Then the dried gel was stained with Ethidium Bromide (Sigma-Aldrich, 2375) and imaged with BioRad GelDocXR imager (BioRad). Then denatured in-gel hybridization was performed. Briefly, the gel was denatured with 0.5 M NaOH + 1.5 M NaCl for 30 min; neutralized with 0.5 M Tris–HCl pH 8.0 + 1.5 M NaCl for 30 minutes; then prehybridized with 10× Denhardt’s buffer + 5× SSC + 0.5% SDS for 30 minutes and hybridized overnight at 42°C with ^32^P-labeled telomeric C-probe prepared as prepared before^6969^. The hybridized gel was washed with 2× SSC + 0.5% SDS and 2× SSC + 0.1% SDS for 3 times each, then exposed to PhosphorImager screen (GE Healthcare) and scanned on Typhoon imager with ImageQuant (Molecular Dynamics).

### Analysis of Telomere-C coverage in the proximal regions

We computed normalized Telomere-C coverages within our loci of interest, extending 3kb into the flanking regions. We analyzed 20,192 protein-coding genes, 388 ITS, 202 TAR1, and 329 D20S16 loci. The collective coverages were converted and scaled into a matrix using the *computeMatrix* function from Deeptools (v3.5.4) and visualized in both a heatmap and a line plot using the *plotHeatmap* function from Deeptools.

### Three-Dimensional Fluorescence in situ hybridization (3D-FISH)

3D-FISH was performed as described previously^22^ with modifications. Cells were cultured to 70% confluency, harvested with 0.25% Trypsin, pelleted by centrifugation at 1,500 rpm for 5 minutes, and resuspended with 0.075 M KCl for 10 minutes. Ice-cold fixative solution (methanol and glacial acetic acid in the ratio of 3:1) was used to fix the cells prior to slide preparation for interphase FISH. 30 µL of cell suspension were dropped on a pre-cleaned slide and hybridized to pan-telomere PNA probe-Cy3 (PNA Bio, F1002) co-hybridized with DNA oligo probes recognizing ITS at chr19:19,306,848-19,307,366 (CHM13v2) (Empire Genomics, RP11-837J10, Green-dUTP), D20S16 arrays at chr20: 49,889,302-49,895,503 (Empire Genomics, RP11-737E14, Green-dUTP) and chr20: 50,241,413-50,286,275 (Empire Genomics, RP11-668L3, Green-dUTP). Slides were denatured at 80°C for 3 minutes followed by hybridization at 37°C overnight. After hybridization, slides were washed with solution 1 (70% formamide solution containing 10 mM Tris-HCl, pH 7.5) twice for 5 minutes followed by a wash with solution 2 (100 mM Tris-HCl, 150 mM NaCl, and 0.08% Tween-20) three times for 5 minutes. Ethanol dehydration was carried out for 5 minutes in 70%, 95%, and 100% ethanol respectively. After drying, the slide was mounted with DAPI Antifade (MetaSystems, D-0902-500-DA) and incubated overnight for hardening. Images were acquired on Nikon Eclipse Ti-2 for Telomere-D20S16 or Zeiss Confocal LSM 880 microscope for Telomere-ITS19 using DAPI, Cy3, and FAM channels. 3D rendering was carried out using the Imaris software (v10.2.0, Oxford Instruments).

### Image analysis

For measurement analysis using Imaris (v10.2.0), at least 30 nuclei were imaged per slide. Image processing was carried out by adjusting the Gaussian filter to 0.107 µm, and background subtraction at a filter width of 61.8 µm. A nucleus was counted as a positive event of co-localization if at least one pair of overlapping foci was identified by manual examination. For spatial analysis, the coordinates of all foci were centered to the corresponding centroid of the nucleus to obtain the normalized coordinates. These coordinates were used to determine the spatial distribution of the foci (both green and red channels) via bivariate Ripley’s K-function with a pooling strategy^70,71^. The analysis was performed using our in-house R script with the *spatstat* package^72^. The bivariate K-function was first calculated for each nucleus, and all the bivariate K-functions were then pooled. The median K-function was used as the observed curve, and the minimum and maximum K-function defined the confidence envelope. We tested for complete spatial randomness using a Poisson distribution as our null hypothesis. The aggregation was determined by comparing the observed curve to the theoretical distribution (an observed curve above the theoretical curve). The p-value and D-statistic (the maximum difference between the observed and theoretical cumulative distributions) were computed by Kolmogorov–Smirnov test.

## Data Availability

Project data has been deposited at https://www.synapse.org/Synapse:syn64463432. Code needed to replicate the analyses performed in this manuscript can be accessed at https://github.com/barthel-lab/Telomere-C and https://github.com/barthel-lab/micro-C_data_processing.

## Author Contributions

F.P.B. and Y.C. conceived the project. Y.C., O.M. and N.H.F. performed the experiments. T.Z. performed the Southern blotting. Y.C. and T.R.R.B. performed the data analysis. Y.C. and F.P.B wrote the manuscript and all authors reviewed the manuscript and provided input.

## Acknowledgements

We would like to acknowledge the Forbeck Foundation for hosting outstanding discussion forums that helped mature the concepts leading to this work. We acknowledge all Barthel lab members for insightful comments that helped elevate the level of this work. This research includes work performed in the TGen Collaborative Sequencing Center, a City of Hope Comprehensive Cancer Center supported shared resource (NCI-P30CA033572). This research was supported by The National Cancer Institutes of the National Institutes of Health under award number R00CA226387.

---

**Supplementary Figure 1.**
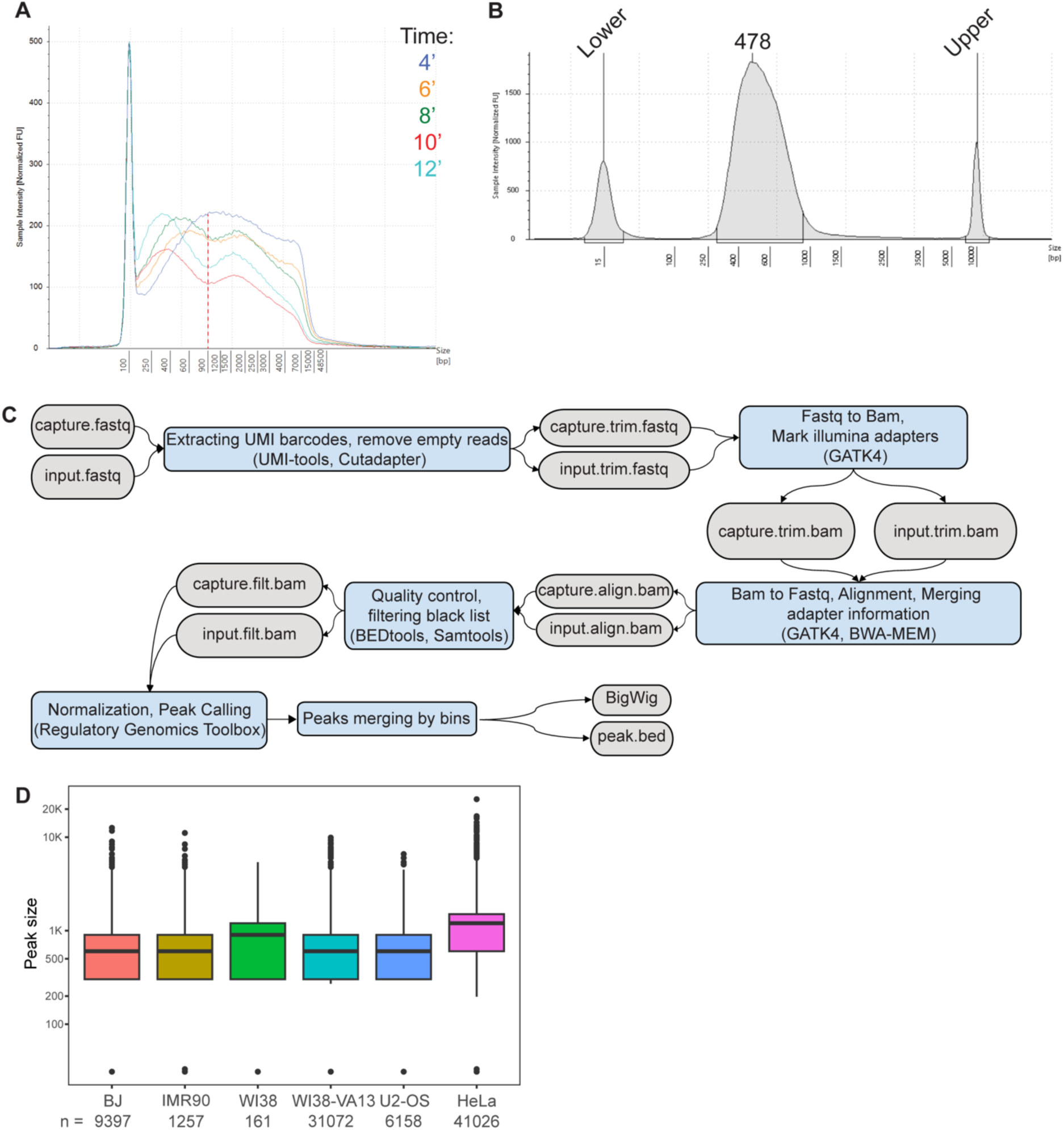
Quality control of Telomere-C library preparation and sequencing data processing. **A** Size distribution of chromatin fragments generated by sonication at indicated durations (4, 6, 8, 10, and 12 minutes). **B** Size distribution of the Telomere-C library, with the peak fragment size indicated. **C** Workflow of Telomere-C data processing, with inputs, intermediate data, and outputs presented in grey boxes; and processing steps and tools shown in blue boxes. **D** Box plot showing the size distribution of Telomere-C peaks across the six cell lines.

**Supplementary Figure 2.**
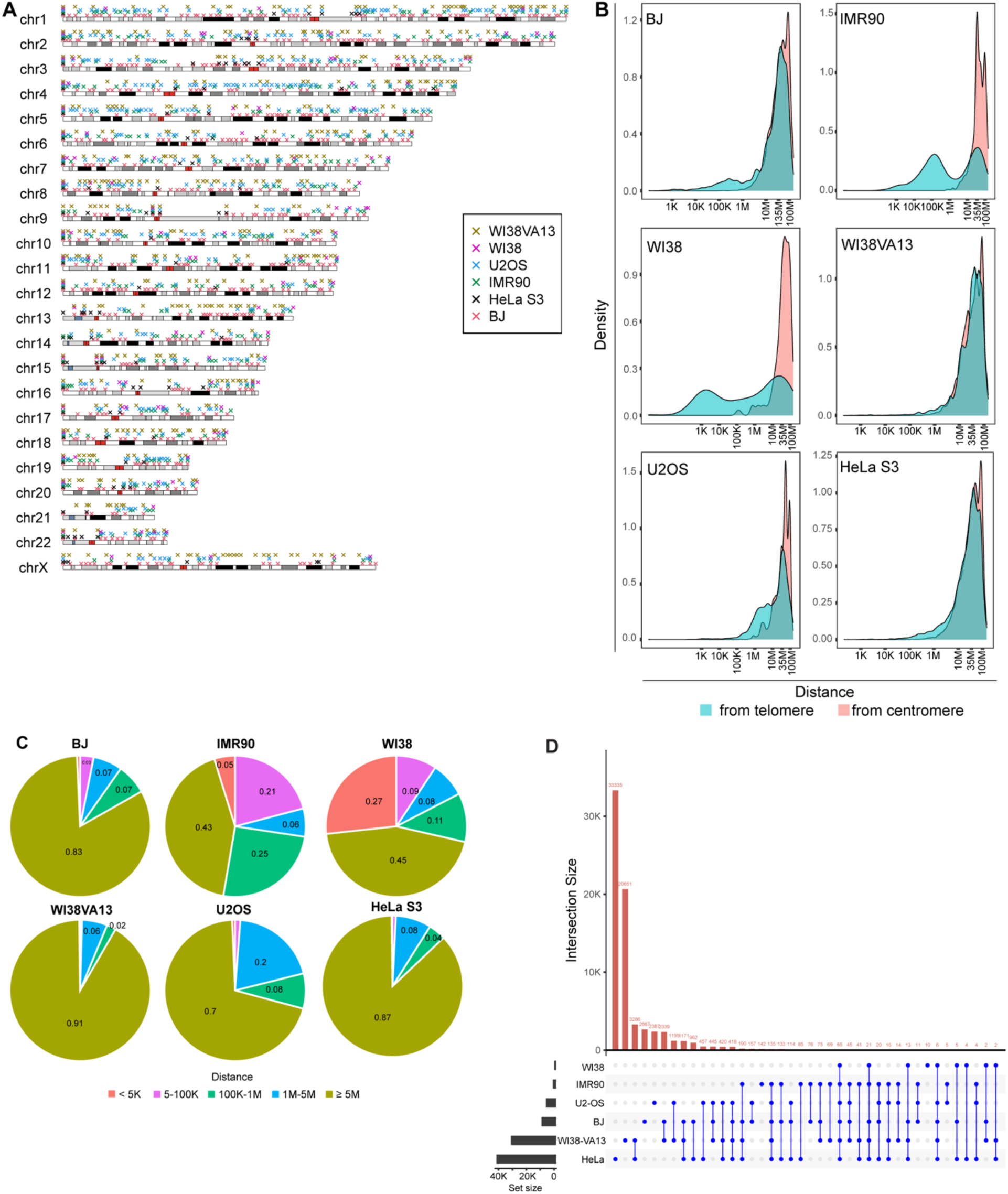
Distribution of Telomere-C peaks on chromosomes. **A** Visualization of Telomere-C peaks (colored crosses) across all chromosomes in BJ (red), HeLa S3 (black), IMR90 (green), U2-OS (blue), WI38 (magenta), and WI38-VA13 (olive). **B** Distribution of the relative distances of Telomere-C peaks to telomeres (cyan) or centromeres (coral) in the indicated cells. **C** Proportion of Telomere-C peak distances to telomeres. Distances are categorized by range: <5 Kb (coral), 5-100 Kb (purple), 100 Kb-1 Mb (blue), 1-5 Mb (green), and ≥ 5 Mb (olive) in the indicated cells. **D** An UpSet plot showing unique and shared Telomere-C peaks identified among the six cell lines.

**Supplementary Figure 3.**
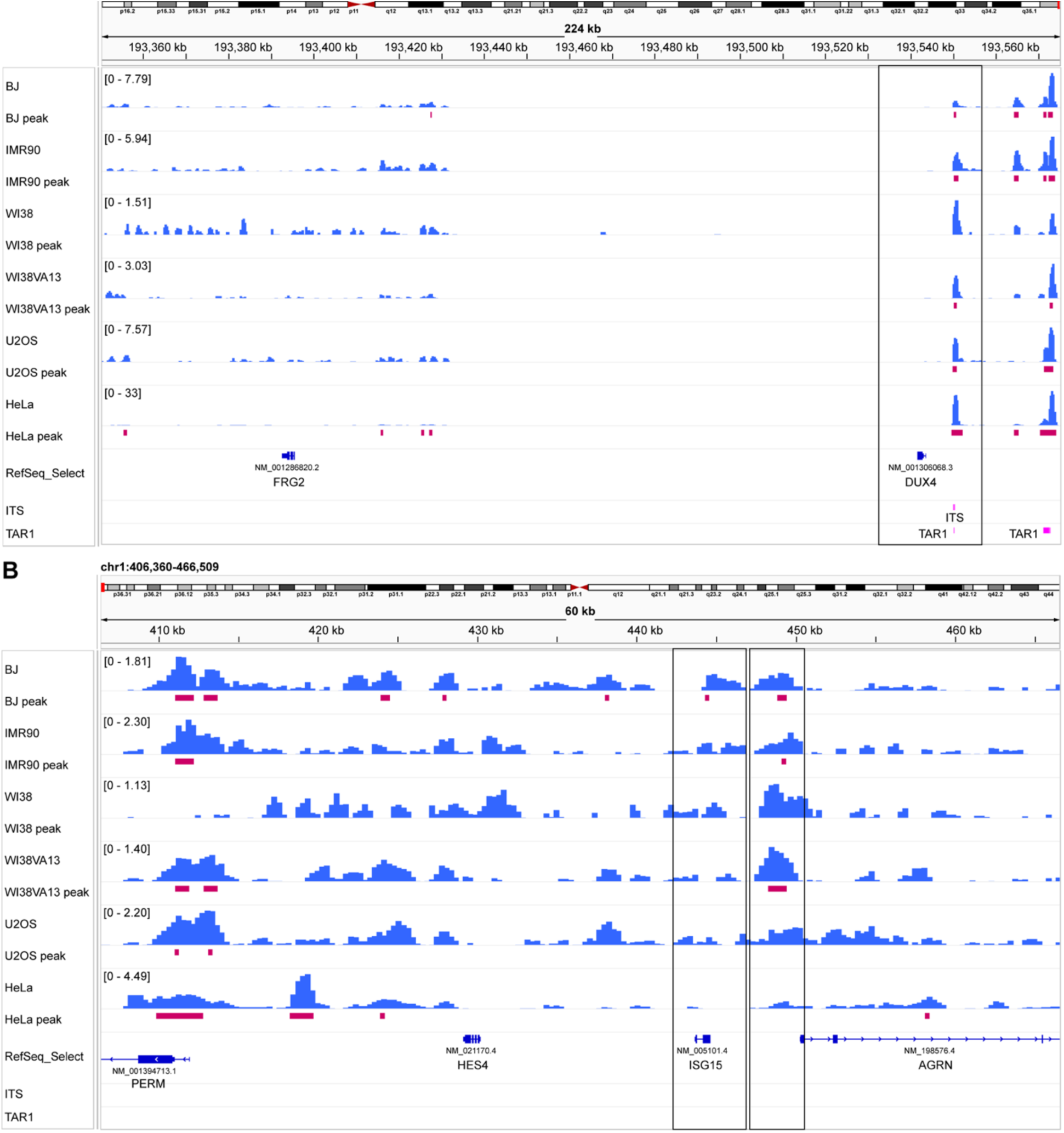
Validation of telomeric chromatin interactions at *DUX4* and *ISG15*. Panels show Telomere-C signals (blue bars) and corresponding called peaks (red rectangles) for each indicated cell line at the **A** *DUX4* and **B** *ISG15* loci. The black rectangles highlight the corresponding transcript and downstream region.

**Supplementary Figure 4.**
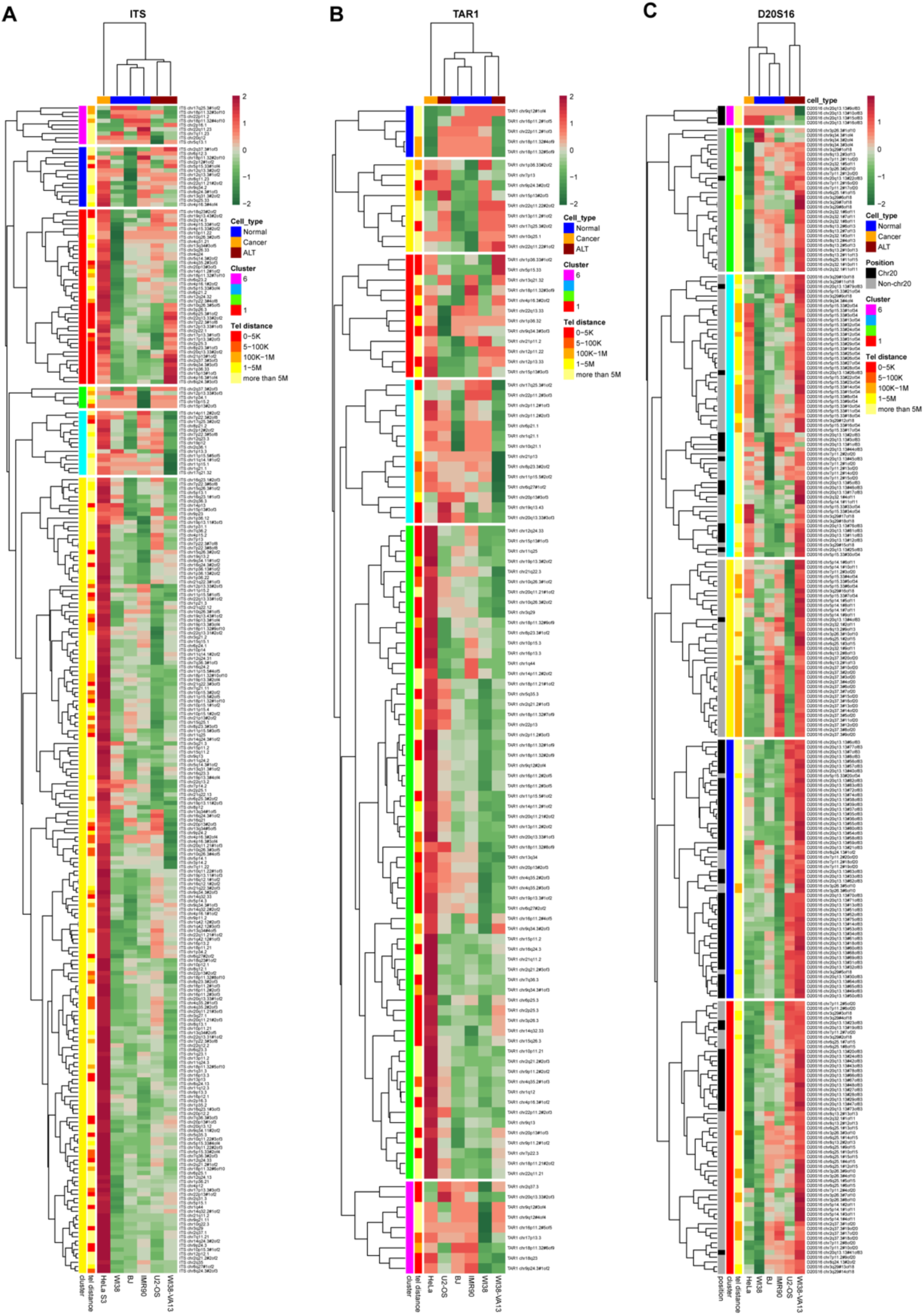
Heatmaps of normalized Telomere-C signals (Z-score) across the repeat elements. The Telomere-C signal intensity is color-coded from green (low, -2) to red (high, 2). Elements (rows) and cell lines (columns) are hierarchically clustered by Euclidean distance. **A** ITS (n = 338); **B** TAR1 (n = 202); and **C** D20S16 (n = 329). Two colored sidebars on the left indicate the assigned cluster of elements (1-6, color-coded: red, yellow, green, cyan, blue, and purple) and the distance to the telomere. A colored-bar at the top indicates cell type: normal (blue), cancer (yellow), and ALT+ (dark red).

**Supplementary Figure 5.**
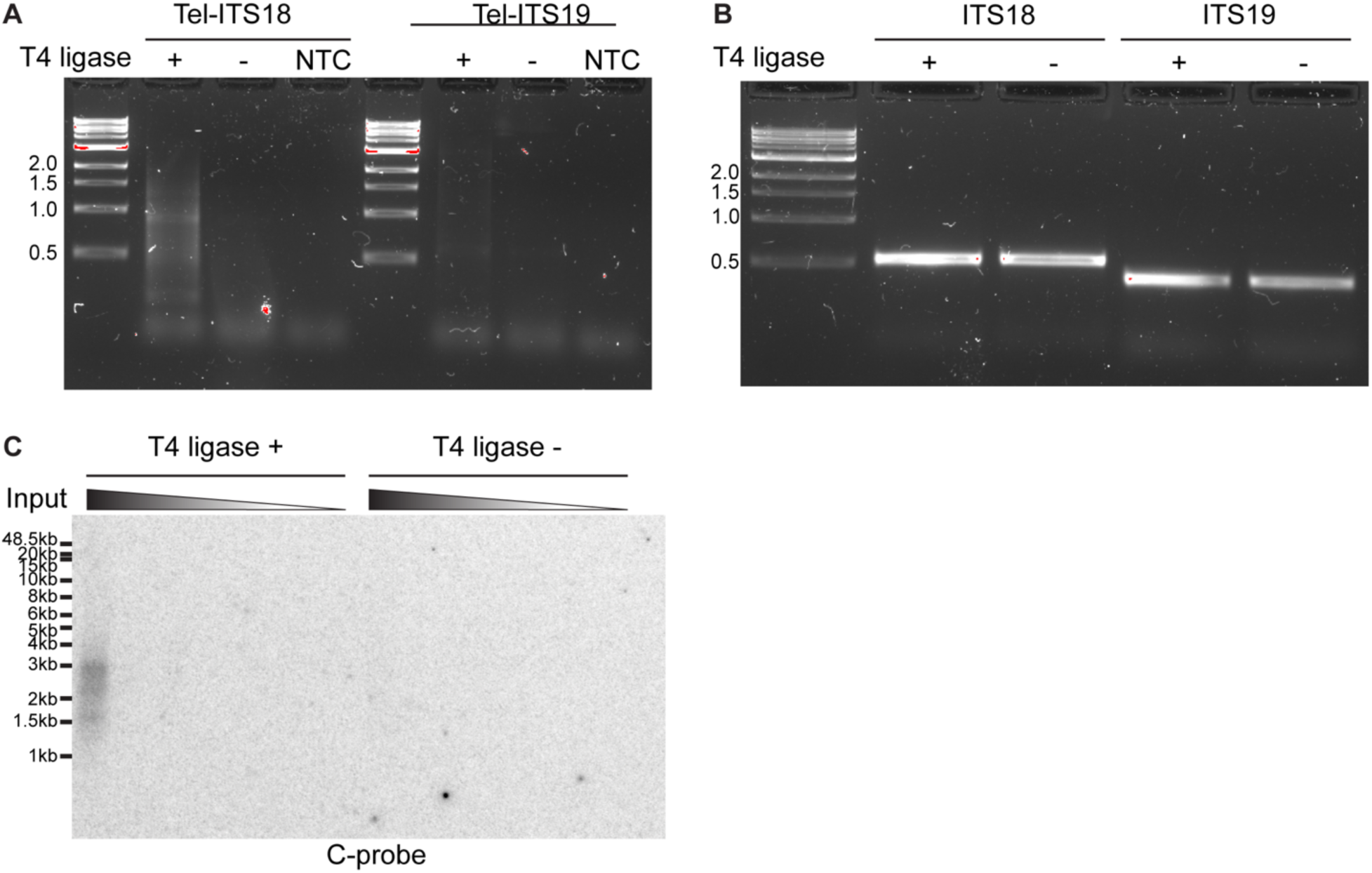
Further validation of Telomere-ITS interactions. **A** PCR results showing the Telomere-ITS18 and Telomere-ITS19 interactions, visualized by electrophoresis on a 1% agarose gel. 3C DNA was used as a template, with or without T4 ligase (proximity ligation). Non-template controls are indicated as NTC. **B** PCR results showing the amplification of the ITS loci themselves (non-ligated control) visualized by electrophoresis on a 1% agarose gel. **C** Visualization of the Telomere-ITS19 amplicon by in-gel hybridization. The amount of DNA template is indicated above the gel. The results were visualized using ^32^P-labeled C-rich telomeric probe.

**Supplementary Figure 6.**
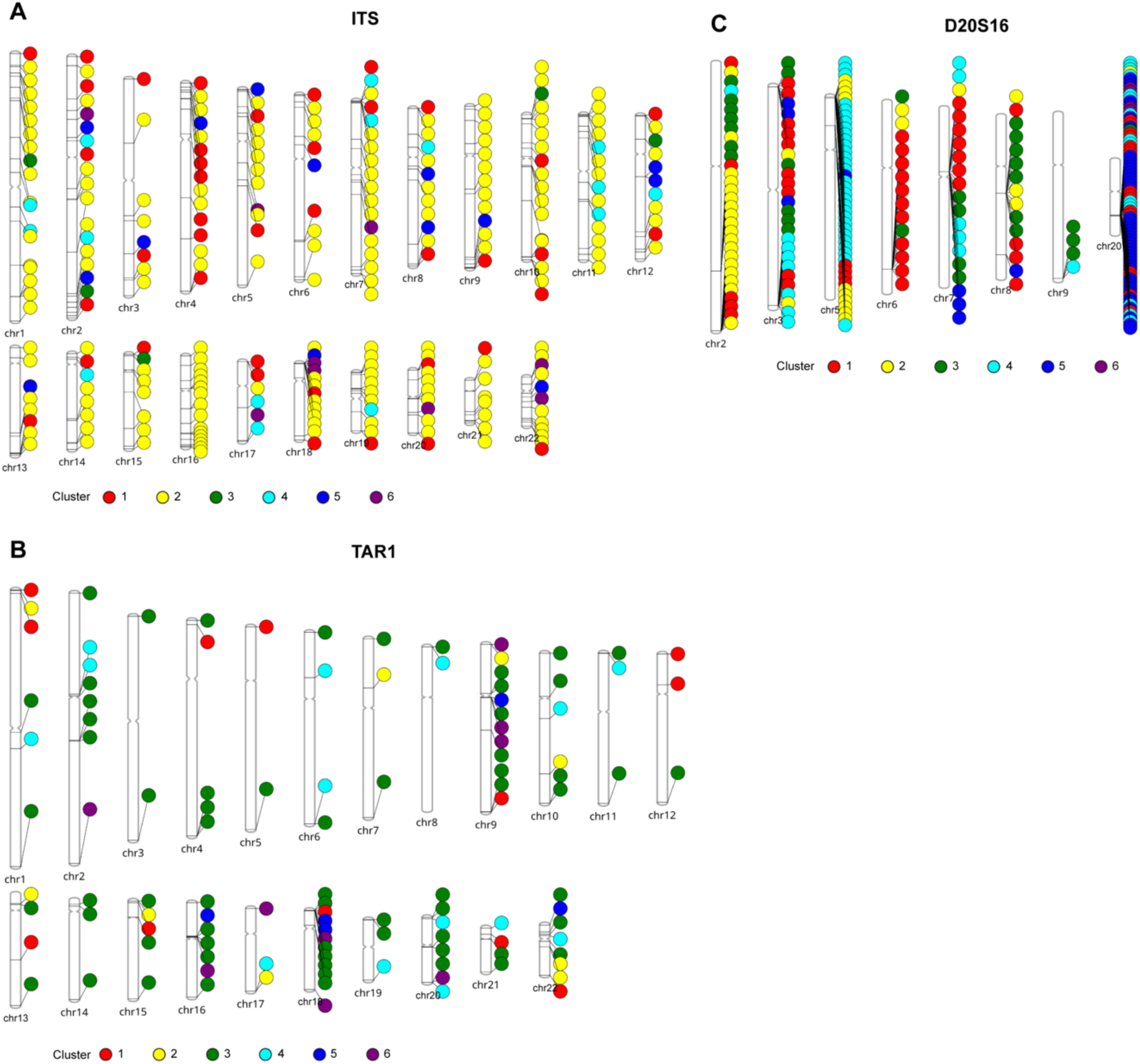
Genomic distribution of Telomere-C associated repetitive elements. PhenoGram showing the distribution of **A** ITS (n = 338); **B** TAR1 (n = 202); and **C** D20S16 (n = 329). Data points are color-coded by their assigned cluster (1-6), where the colors correspond to red, yellow, green, cyan, blue, and purple. Chromosomes without data points are omitted from the visualization.

**Supplementary Figure 7.**
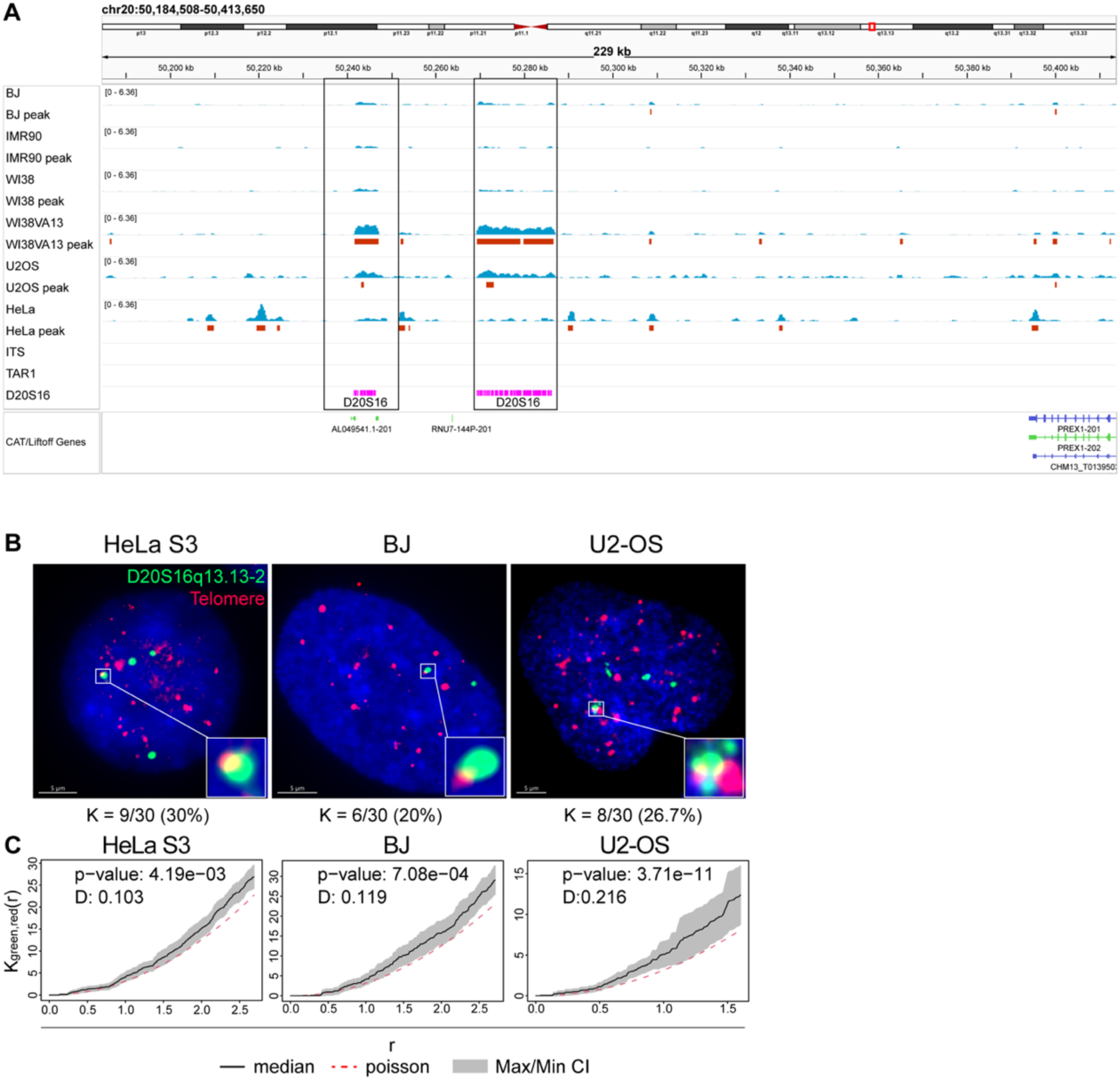
Validation of Telomere-D20S16 interactions at chr20q13. **A** IGV screenshot showing Telomere-C signals (blue bars), corresponding called peaks (red rectangles) and annotated repeat elements (magenta rectangles) for each indicated cell line at the *D20S16* locus (chr20q13.13). The black rectangles highlight the corresponding regions of interest: *left* chr20q13.13-2; *right* chr20q13.13. **B** 3D-FISH images with scale bars (5 µm) demonstrating the colocalization between D20S16 (chr20q13.13-2, green) and telomeres (red) in HeLa S3, BJ, and U2-OS cells. Nuclei were counterstained with DAPI (blue). The enlarged window represents a 5X magnification of selected colocalization foci. The number and percentage of colocalized nuclei are indicated below. **C** Pooled Ripley’s K-function plot illustrating the aggregation levels between D20S16 and telomeres in HeLa S3, BJ, and U2-OS cells. The solid back line indicates the median of the bivariate K-function, with the grey shaded area indicating the confidence intervals. The red dotted line indicates the theoretical Poisson distribution (null hypothesis). The p-value and D statistics were calculated using Kolmogorov– Smirnov test.

## References

1 Decker, M. L., Chavez, E., Vulto, I. & Lansdorp, P. M. Telomere length in Hutchinson-Gilford Progeria Syndrome. Mechanisms of Ageing and Development 130, 377–383, doi:10.1016/j.mad.2009.03.001 (2009).

2 Wood, A. M. et al. TRF2 and lamin A/C interact to facilitate the functional organization of chromosome ends. Nature Communications 5, 5467, doi:10.1038/ncomms6467 (2014).

3 Moyzis, R. K. et al. A highly conserved repetitive DNA sequence, (TTAGGG)n, present at the telomeres of human chromosomes. Proceedings of the National Academy of Sciences of the United States of America 85, 6622–6626, doi:10.1073/pnas.85.18.6622 (1988).

4 Okuda, K. et al. Telomere length in the newborn. Pediatr Res 52, 377–381, doi:10.1203/00006450-200209000-00012 (2002).

5 Canela, A., Vera, E., Klatt, P. & Blasco, M. A. High-throughput telomere length quantification by FISH and its application to human population studies. Proceedings of the National Academy of Sciences of the United States of America 104, 5300–5305, doi:10.1073/pnas.0609367104 (2007).

6 Makarov, V. L., Hirose, Y. & Langmore, J. P. Long G Tails at Both Ends of Human Chromosomes Suggest a C Strand Degradation Mechanism for Telomere Shortening. Cell 88, 657–666, doi:10.1016/S0092-8674(00)81908-X (1997).

7 de Lange, T. How telomeres solve the end-protection problem. Science 326, 948–952, doi:10.1126/science.1170633 (2009).

8 Stansel, R. M., de Lange, T. & Griffith, J. D. T-loop assembly in vitro involves binding of TRF2 near the 3’ telomeric overhang. The EMBO journal 20, 5532–5540, doi:10.1093/emboj/20.19.5532 (2001).

9 Amiard, S. et al. A topological mechanism for TRF2-enhanced strand invasion. Nature Structural & Molecular Biology 14, 147–154, doi:10.1038/nsmb1192 (2007).

10 Griffith, J. D. et al. Mammalian telomeres end in a large duplex loop. Cell 97, 503–514, doi:10.1016/s0092-8674(00)80760-6 (1999).

11 van Steensel, B., Smogorzewska, A. & de Lange, T. TRF2 protects human telomeres from end-to-end fusions. Cell 92, 401–413, doi:10.1016/s0092-8674(00)80932-0 (1998).

12 Wang, R. C., Smogorzewska, A. & de Lange, T. Homologous recombination generates T-loop-sized deletions at human telomeres. Cell 119, 355–368, doi:10.1016/j.cell.2004.10.011 (2004).

13 Gottschling, D. E., Aparicio, O. M., Billington, B. L. & Zakian, V. A. Position effect at S. cerevisiae telomeres: Reversible repression of Pol II transcription. Cell 63, 751–762, 10.1016/0092-8674(90)90141-Z (1990).

14 de Bruin, D., Zaman, Z., Liberatore, R. A. & Ptashne, M. Telomere looping permits gene activation by a downstream UAS in yeast. Nature 409, 109–113, doi:10.1038/35051119 (2001).

15 Stavenhagen, J. B. & Zakian, V. A. Yeast telomeres exert a position effect on recombination between internal tracts of yeast telomeric DNA. Genes Dev. 12, 3044–3058, doi:10.1101/gad.12.19.3044 (1998).

16 Tham, W. H. & Zakian, V. A. Transcriptional silencing at Saccharomyces telomeres: implications for other organisms. Oncogene 21, 512–521, doi:10.1038/sj.onc.1205078 (2002).

17 Strahl-Bolsinger, S., Hecht, A., Luo, K. & Grunstein, M. SIR2 and SIR4 interactions differ in core and extended telomeric heterochromatin in yeast. Genes Dev. 11, 83–93, doi:10.1101/gad.11.1.83 (1997).

18 Bystricky, K., Laroche, T., van Houwe, G., Blaszczyk, M. & Gasser, S. M. Chromosome looping in yeast: telomere pairing and coordinated movement reflect anchoring efficiency and territorial organization. The Journal of Cell Biology 168, 375–387, doi:10.1083/jcb.200409091 (2005).

19 Miele, A., Bystricky, K. & Dekker, J. Yeast silent mating type loci form heterochromatic clusters through silencer protein-dependent long-range interactions. PLoS genetics 5, e1000478, doi:10.1371/journal.pgen.1000478 (2009).

20 Lou, Z. et al. Telomere length regulates ISG15 expression in human cells. Aging (Albany NY) 1, 608–621, doi:10.18632/aging.100066 (2009).

21 Robin, J. D. et al. SORBS2 transcription is activated by telomere position effect-over long distance upon telomere shortening in muscle cells from patients with facioscapulohumeral dystrophy. Genome Res. 25, 1781–1790, doi:10.1101/gr.190660.115 (2015).

22 Kim, W. et al. Regulation of the Human Telomerase Gene TERT by Telomere Position Effect—Over Long Distances (TPE-OLD): Implications for Aging and Cancer. PLOS Biology 14, e2000016, doi:10.1371/journal.pbio.2000016 (2016).

23 Lafontaine, D. L., Yang, L., Dekker, J. & Gibcus, J. H. Hi-C 3.0: Improved Protocol for Genome-Wide Chromosome Conformation Capture. Curr Protoc 1, e198, doi:10.1002/cpz1.198 (2021).

24 Stadler, G. et al. Telomere position effect regulates DUX4 in human facioscapulohumeral muscular dystrophy. Nat Struct Mol Biol 20, 671–678, doi:10.1038/nsmb.2571 (2013).

25 Robin, J. D. et al. Telomere position effect: regulation of gene expression with progressive telomere shortening over long distances. Genes Dev. 28, 2464–2476, doi:10.1101/gad.251041.114 (2014).

26 Kim, N. W. et al. Specific association of human telomerase activity with immortal cells and cancer. Science 266, 2011–2015, doi:10.1126/science.7605428 (1994).

27 Bryan, T. M., Englezou, A., Dalla-Pozza, L., Dunham, M. A. & Reddel, R. R. Evidence for an alternative mechanism for maintaining telomere length in human tumors and tumor-derived cell lines. Nat Med 3, 1271–1274, doi:10.1038/nm1197-1271 (1997).

28 Zhang, J. M., Yadav, T., Ouyang, J., Lan, L. & Zou, L. Alternative Lengthening of Telomeres through Two Distinct Break-Induced Replication Pathways. Cell Rep 26, 955–968 e953, doi:10.1016/j.celrep.2018.12.102 (2019).

29 Dilley, R. L. et al. Break-induced telomere synthesis underlies alternative telomere maintenance. Nature 539, 54–58, doi:10.1038/nature20099 (2016).

30 Conomos, D., Reddel, R. R. & Pickett, H. A. NuRD-ZNF827 recruitment to telomeres creates a molecular scaffold for homologous recombination. Nat Struct Mol Biol 21, 760–770, doi:10.1038/nsmb.2877 (2014).

31 Roukos, V. et al. Spatial dynamics of chromosome translocations in living cells. Science 341, 660–664, doi:10.1126/science.1237150 (2013).

32 Dejardin, J. & Kingston, R. E. Purification of proteins associated with specific genomic Loci. Cell 136, 175–186, doi:10.1016/j.cell.2008.11.045 (2009).

33 Conomos, D. et al. Variant repeats are interspersed throughout the telomeres and recruit nuclear receptors in ALT cells. J Cell Biol 199, 893–906, doi:10.1083/jcb.201207189 (2012).

34 Marzec, P. et al. Nuclear-receptor-mediated telomere insertion leads to genome instability in ALT cancers. Cell 160, 913–927, doi:10.1016/j.cell.2015.01.044 (2015).

35 Gaela, V. M. et al. Orphan nuclear receptors-induced ALT-associated PML bodies are targets for ALT inhibition. Nucleic Acids Res 52, 6472–6489, doi:10.1093/nar/gkae389 (2024).

36 Kim, W. & Shay, J. W. Long-range telomere regulation of gene expression: Telomere looping and telomere position effect over long distances (TPE-OLD). Differentiation 99, 1–9, doi:10.1016/j.diff.2017.11.005 (2018).

37 Lieberman-Aiden, E. et al. Comprehensive mapping of long-range interactions reveals folding principles of the human genome. *Science (New York*, N.Y.) 326, 289–293, doi:10.1126/science.1181369 (2009).

38 Fullwood, M. J. et al. An oestrogen-receptor-alpha-bound human chromatin interactome. Nature 462, 58–64, doi:10.1038/nature08497 (2009).

39 Mumbach, M. R. et al. HiChIP: efficient and sensitive analysis of protein-directed genome architecture. Nature Methods 13, 919–922, doi:10.1038/nmeth.3999 (2016).

40 Kimura, M. et al. Measurement of telomere length by the Southern blot analysis of terminal restriction fragment lengths. Nature Protocols 5, 1596–1607, doi:10.1038/nprot.2010.124 (2010).

41 Hsieh, T. H. et al. Mapping Nucleosome Resolution Chromosome Folding in Yeast by Micro-C. Cell 162, 108–119, doi:10.1016/j.cell.2015.05.048 (2015).

42 Cawthon, R. M. Telomere measurement by quantitative PCR. Nucleic Acids Research 30, 47e–47, doi:10.1093/nar/30.10.e47 (2002).

43 Lai, T. P. et al. A method for measuring the distribution of the shortest telomeres in cells and tissues. Nat Commun 8, 1356, doi:10.1038/s41467-017-01291-z (2017).

44 Hazan, J., Dubay, C., Pankowiak, M. P., Becuwe, N. & Weissenbach, J. A genetic linkage map of human chromosome 20 composed entirely of microsatellite markers. Genomics 12, 183–189, doi:10.1016/0888-7543(92)90364-x (1992).

45 Yandim, C. & Karakulah, G. Dysregulated expression of repetitive DNA in ER+/HER2-breast cancer. Cancer Genet 239, 36–45, doi:10.1016/j.cancergen.2019.09.002 (2019).

46 Yandim, C. & Karakulah, G. Expression dynamics of repetitive DNA in early human embryonic development. BMC Genomics 20, 439, doi:10.1186/s12864-019-5803-1 (2019).

47 Hu, Y., Mizuguchi, K. & Hashimoto, K. Unveiling unique expression patterns of D20S16 satellite DNA in human embryonic development. Scientific Reports 15, 26770, doi:10.1038/s41598-025-11753-w (2025).

48 Yang, D. et al. Human telomeric proteins occupy selective interstitial sites. Cell Research 21, 1013–1027, doi:10.1038/cr.2011.39 (2011).

49 Simonet, T. et al. The human TTAGGG repeat factors 1 and 2 bind to a subset of interstitial telomeric sequences and satellite repeats. Cell Research 21, 1028–1038, doi:10.1038/cr.2011.40 (2011).

50 Arnoult, N., Van Beneden, A. & Decottignies, A. Telomere length regulates TERRA levels through increased trimethylation of telomeric H3K9 and HP1α. Nature Structural & Molecular Biology 19, 948–956, doi:10.1038/nsmb.2364 (2012).

51 Fernandes, R. V., Feretzaki, M. & Lingner, J. The makings of TERRA R-loops at chromosome ends. Cell Cycle 20, 1745–1759, doi:10.1080/15384101.2021.1962638 (2021).

52 Mokhtaridoost, M. et al. Inter-chromosomal contacts demarcate genome topology along a spatial gradient. Nature Communications 15, 9813, doi:10.1038/s41467-024-53983-y (2024).

53 Smith, T., Heger, A. & Sudbery, I. UMI-tools: modeling sequencing errors in Unique Molecular Identifiers to improve quantification accuracy. Genome Res. 27, 491–499, doi:10.1101/gr.209601.116 (2017).

54 Martin, M. Cutadapt removes adapter sequences from high-throughput sequencing reads. EMBnet.journal 17, 10–12, doi:10.14806/ej.17.1.200 (2011).

55 Auwera, G. V. d. & O’Connor, B. D. Genomics in the cloud: using Docker, GATK, and WDL in Terra. First edition edn, (O’Reilly, 2020).

56 Rhie, A. et al. The complete sequence of a human Y chromosome. Nature 621, 344–354, doi:10.1038/s41586-023-06457-y (2023).

57 Li, H. Aligning sequence reads, clone sequences and assembly contigs with BWA-MEM. arXiv: Genomics (2013).

58 Ogata, J. D. et al. excluderanges: exclusion sets for T2T-CHM13, GRCm39, and other genome assemblies. Bioinformatics 39, btad198, doi:10.1093/bioinformatics/btad198 (2023).

59 Quinlan, A. R. & Hall, I. M. BEDTools: a flexible suite of utilities for comparing genomic features. Bioinformatics 26, 841–842, doi:10.1093/bioinformatics/btq033 (2010).

60 Li, H. et al. The Sequence Alignment/Map format and SAMtools. *Bioinformatics (Oxford*, England) 25, 2078–2079, doi:10.1093/bioinformatics/btp352 (2009).

61 Li, Z. et al. RGT: a toolbox for the integrative analysis of high throughput regulatory genomics data. BMC Bioinformatics 24, 79, doi:10.1186/s12859-023-05184-5 (2023).

62 Gel, B. & Serra, E. karyoploteR: an R/Bioconductor package to plot customizable genomes displaying arbitrary data. Bioinformatics 33, 3088–3090, doi:10.1093/bioinformatics/btx346 (2017).

63 Robinson, J. T. et al. Integrative genomics viewer. Nature Biotechnology 29, 24–26, doi:10.1038/nbt.1754 (2011).

64 Conway, J. R., Lex, A. & Gehlenborg, N. UpSetR: an R package for the visualization of intersecting sets and their properties. Bioinformatics 33, 2938–2940, doi:10.1093/bioinformatics/btx364 (2017).

65 Ramírez, F. et al. deepTools2: a next generation web server for deep-sequencing data analysis. Nucleic Acids Research 44, W160–W165, doi:10.1093/nar/gkw257 (2016).

66 Kolde, R. pheatmap: Pretty Heatmaps. (2025).

67 Wolfe, D., Dudek, S., Ritchie, M. D. & Pendergrass, S. A. Visualizing genomic information across chromosomes with PhenoGram. BioData Min 6, 18, doi:10.1186/1756-0381-6-18 (2013).

68 de Wit, E. & de Laat, W. A decade of 3C technologies: insights into nuclear organization. Genes Dev. 26, 11–24, doi:10.1101/gad.179804.111 (2012).

69 Zhao, Y., Shay, J. W. & Wright, W. E. Telomere G-overhang length measurement method 1: the DSN method. Methods Mol Biol 735, 47–54, doi:10.1007/978-1-61779-092-8_5 (2011).

70 Lagache, T., Meas-Yedid, V. & Olivo-Marin, J.-C. in 2013 IEEE 10th International Symposium on Biomedical Imaging. 896–901 (IEEE).

71 Lagache, T., Sauvonnet, N., Danglot, L. & Olivo-Marin, J. C. Statistical analysis of molecule colocalization in bioimaging. Cytometry Part A 87, 568–579 (2015).

72 Baddeley, A., Rubak, E. & Turner, R. Spatial point patterns : methodology and applications with R. (CRC Press, Taylor & Francis Group, 2016).

